# Global historic pandemics caused by the FAM-1 genotype of the Irish potato pathogen *Phytophthora infestans*

**DOI:** 10.1101/2020.08.25.266759

**Authors:** Amanda C. Saville, Jean B. Ristaino

## Abstract

The FAM-1 genotype of *Phytophthora infestans* caused late blight in the 1840s in the US and Europe and was responsible for the Irish famine. We examined 140 herbarium specimens collected between 1845 and 1991 from six continents and used 12-plex microsatellite genotyping (SSR) to identify FAM-1 and the mtDNA lineage (Herb-1/ Ia) present in historic samples. FAM-1 was detected in approximately 73% of the historic specimens and was found on 6 continents. The US-1 genotype was found in only 27% of the samples and was found later on all continents except Australia/Oceania. FAM-1 was the first genotype detected in almost all the former British colonies from which samples were available. The data from historic samples suggest the FAM-1 genotype was widespread, diverse, and spread more widely than US-1. The famine lineage spread to six continents over 140 years, and likely spread during global colonization from Europe.

## Introduction

Emerging plant diseases threaten crop production and forest ecosystems worldwide^1,2^. Movement of pathogens due to increased trade of plants and plant products has exacerbated outbreaks of plant diseases. *Phytophthora infestans* (Mont.) de Bary caused the Irish potato famine of the 1840s^2^, limits potato and tomato production today, and threatens global food security worldwide^1^. While *P. infestans* can be spread aerially through asexual sporangia, long distance movement of the pathogen is mainly due to transport of infected tubers for use as seed potatoes^2^.

The history of *P. infestans* consists of a series of migrations, combined with periodic displacements of one clonal lineage with another, both of which have occurred at local and global scales^(2–5^. The disease first appeared in the US in 1843, near the ports of New York, Philadelphia and in surrounding states (Supplementary Table 1) ^4,5^. The disease was first reported on the European continent in Belgium in 1845, after which it spread throughout Europe and then into Ireland^2,5,6^. Historic populations of the pathogen in the US and Europe have been studied using mycological herbarium specimens to better understand the pathogen’s origin, identify the outbreak strain, and track its spread from the Americas to Europe^(6–13^. Herbarium specimens collected in the 1840s and later from original outbreak specimens revealed that the famine lineage was a Ia mitochondrial haplotype, disputing previous theories that the US-1 (Ib haplotype) lineage caused the famine ^2,6,8,12,13^. The clonal lineage that caused the Irish potato famine (FAM-1) was identified, the genome was sequenced, and it’s shared ancestry with *P. andina*, a sister species from South America, was documented^6,11^.

Microsatellite genotyping (SSRs) has also been used widely to study the population biology of *P. infestans*^2,10,14^. *P. infestans*-infected leaves from historic specimens collected in North America and Europe were genotyped using SSRs and migration from North America into Europe was documented^10^. The FAM-1 genotype caused the first outbreaks in the US and Europe and was eventually displaced by the newly emerging US-1 genotype around the 1930s-1950s^9,10^. US-1 continued to persist globally until the 1980s, when it was replaced by more aggressive lineages out of Mexico and Europe, but can still be found in select populations today^3,15^.

Historic migrations of the pathogen into the US and Europe have been studied, but little is known about migrations of the FAM-1 genotype to other continents after the original 19^th^ century outbreaks^6,10,12,13^. It has been suggested that FAM-1 was less fit and went extinct^8,16^. The earliest known records of *P. infestans* in Asia indicate it was present in India between 1870 and 1880^17^, in Australia, in Tasmania in 1907^4,18^, and in Kenya in 1941^19^. Genotyping of samples of *P. infestans* from eastern Africa from the mid-21^st^ century revealed the presence of the US-1 genotype^20,21^. Recent genotyping of *P. infestans* from herbarium samples from South and Central America identified the FAM-1 genotype in Colombia, Guatemala, and Costa Rica between the 1910s and 1940s, suggesting that the FAM-1 genotype was present in these regions for many years^2,10^.

The goal of this study was to examine the population structure of historic *P. infestans* using a large global set of outbreak samples^22^. The primary objectives of this research were: (1) to infer the population structure of historic *P. infestans* on six continents (2) Examine the spatial biogeography and diversity of FAM-1 and US-1 genotypes over space and time; and (3) compare the impact of host diversity on genotype diversity and (4) infer putative migration pathways of the pathogen into Africa, Asia and Australia/Oceania.

## Results

### Population Structure

A total of 137 historic samples were genotyped with microsatellites (Table 1). The FAM-1 (n=101) and US-1 (n=36) genotypes were identified in the set of historic samples. A subset of 67 samples were genotyped for mtDNA haplotype and identified as the Herb-1 mitochondrial haplotype, while 32 specimens, all US_1’s were the Ib mitochondrial haplotype. The FAM-1 genotype was the first lineage detected in specimens from most countries sampled (Table 1). (Fig. 1)

**Table 1:**
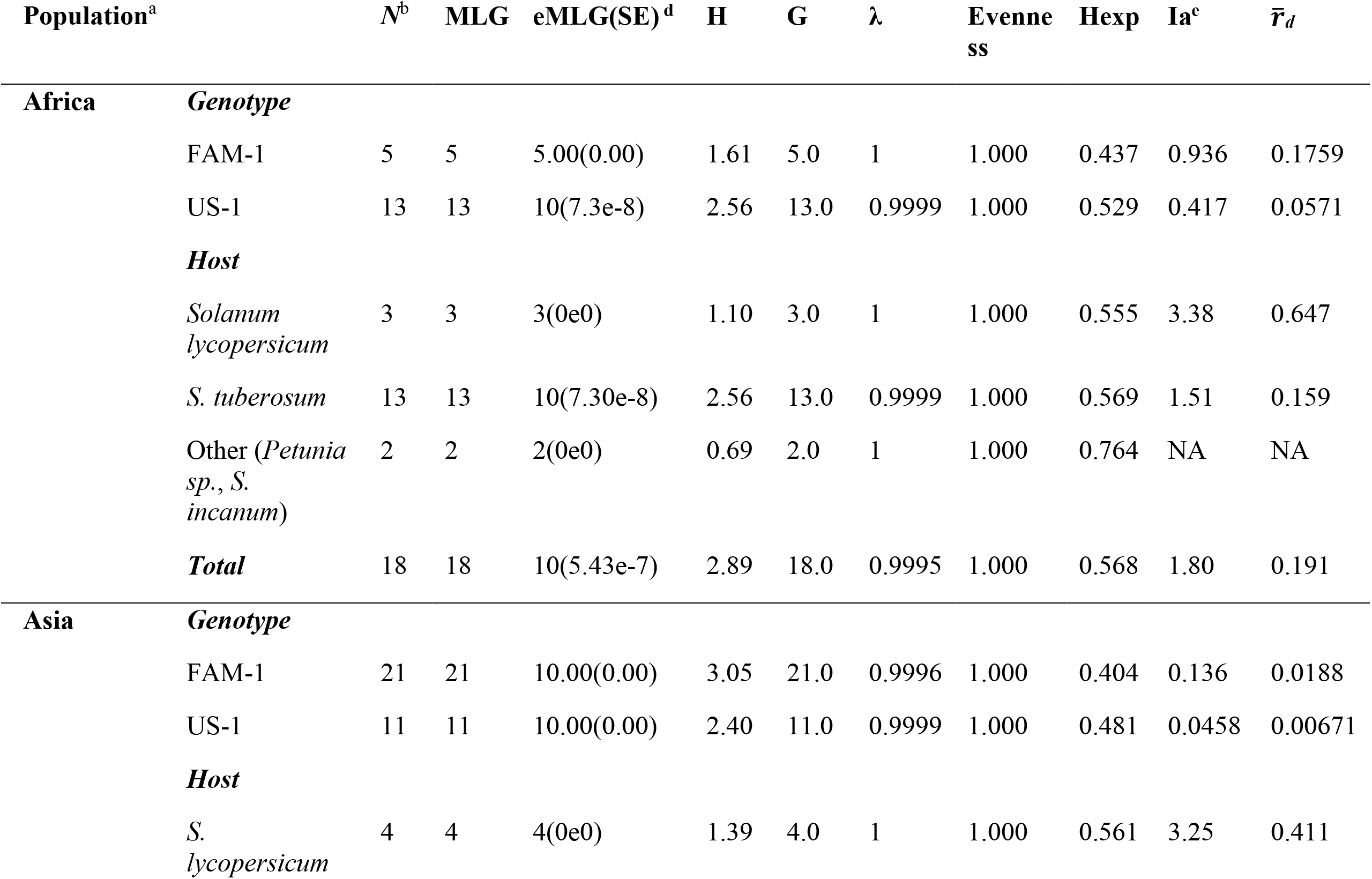
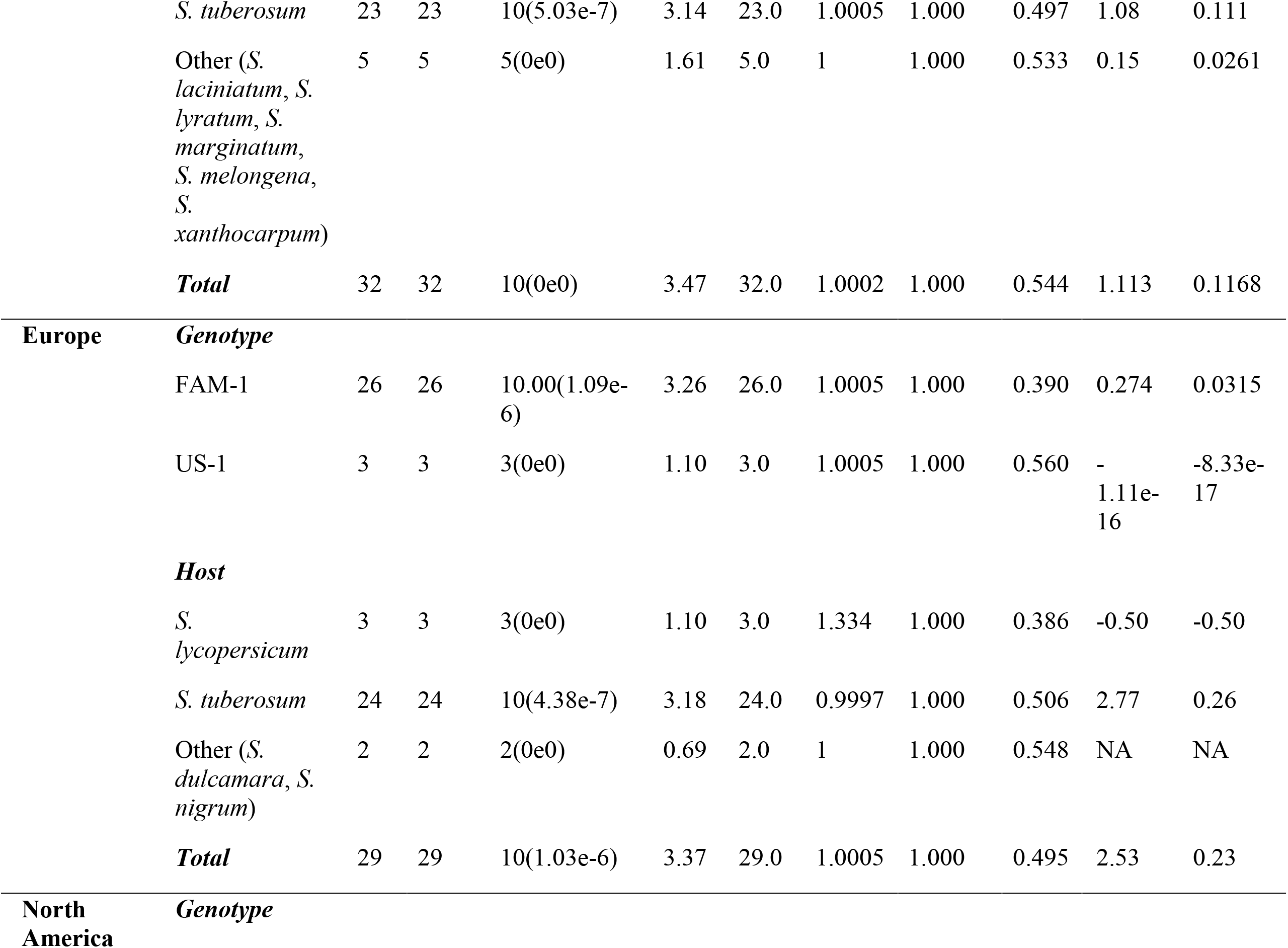
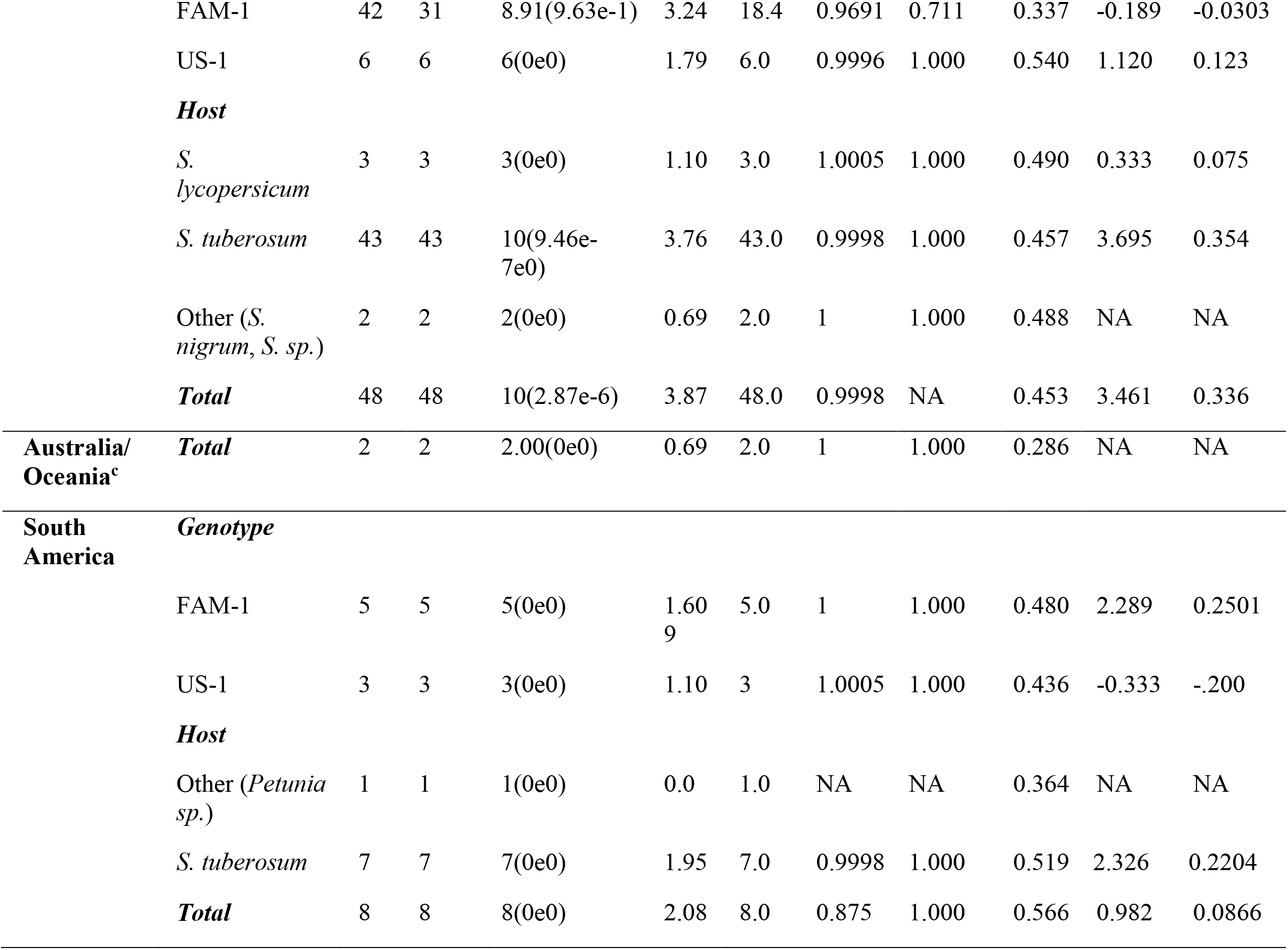
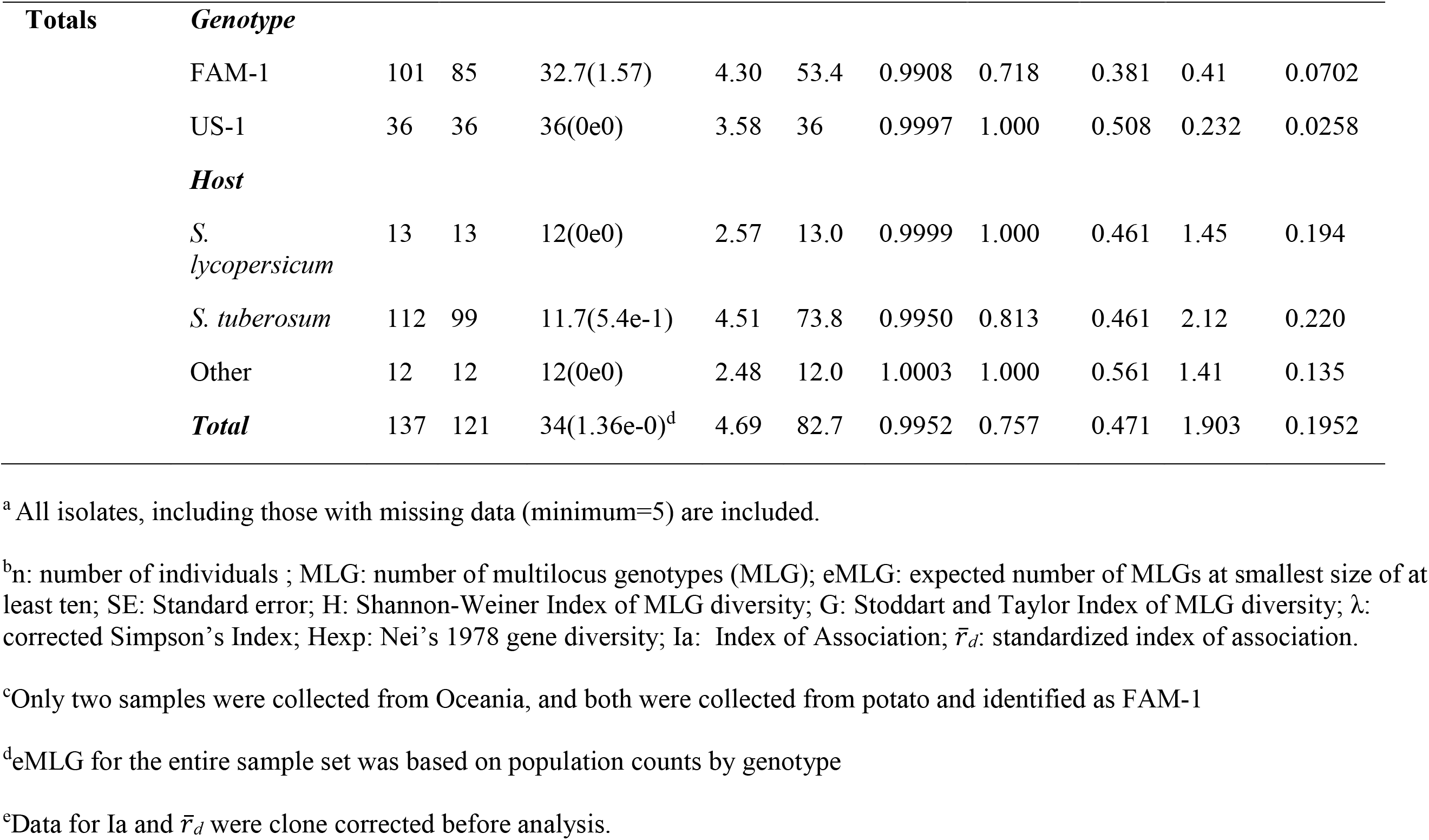
Population statistics of populations of *Phytophthora infestans* from historic global outbreaks sorted by genotype and by region and based on microsatellite data obtained from herbarium specimens.

**Figure 1:**
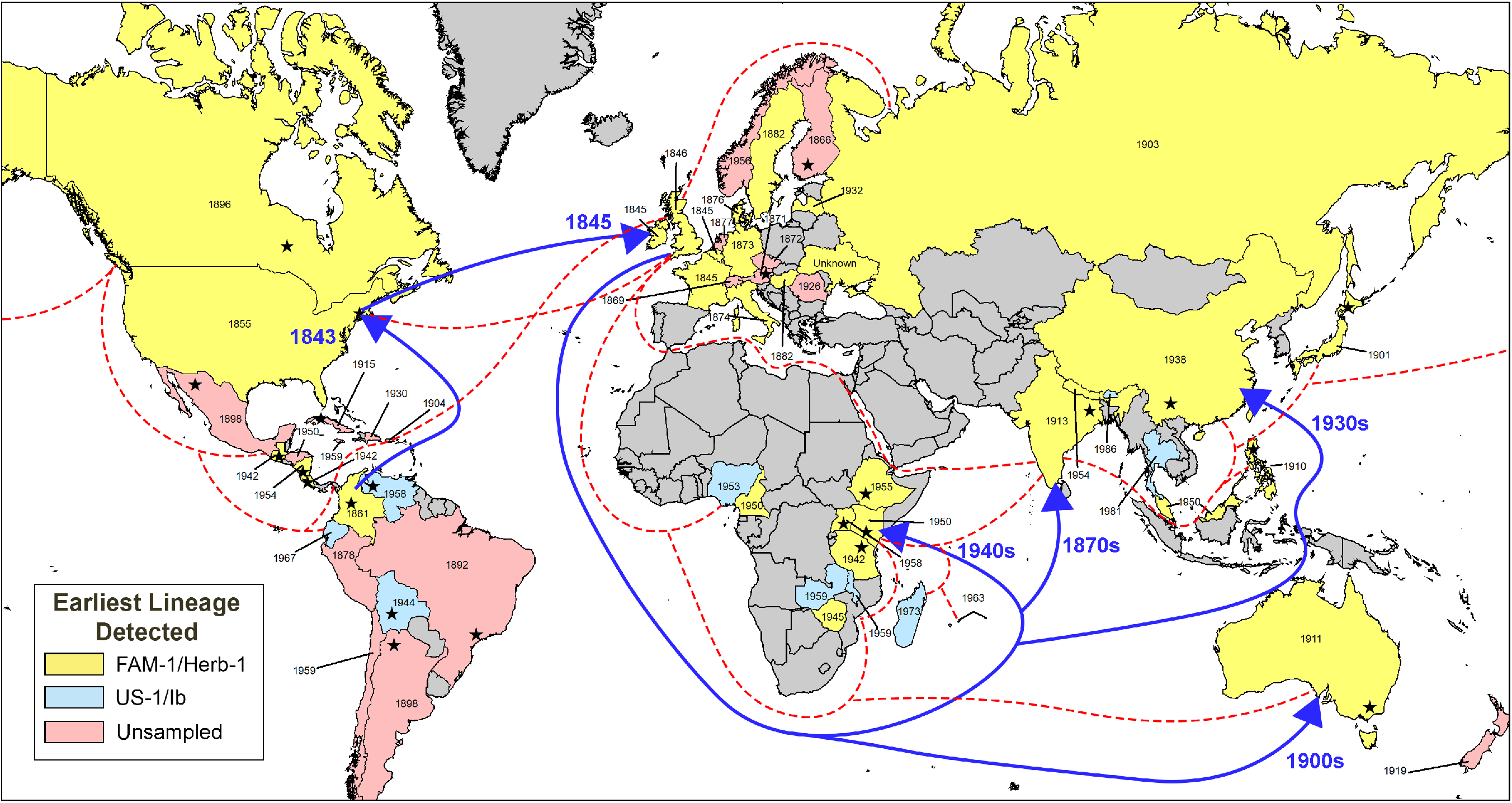
Global map of early outbreaks of late blight caused by *Phytophthora infestans.* Years within each country indicate the date of the earliest known specimen, while color indicates if the genotype was FAM-1, US-1 or unsampled. Dotted lines indicate representative trade routes of the British Empire circa 1932. Arrows indicate the most likely migration path taken by the FAM-1 lineage into Africa and Asia based on DIYABC analysis and trade routes. Stars within each country indicate the approximate location of the first recorded outbreak, if known.

The earliest FAM-1 genotype was found in France in 1845 (K 71) and the most recent FAM-1 was found in Malaysia in 1987 (K 174) (Fig 1, Table1). This indicates that the famine lineage circulated for more than 144 years. In contrast, the earliest US-1 genotype was identified later in the US in 1931 (BPI 186927) and the most recent sample was from India in 1991 (IMI 344673) indicating that US-1 genotype circulated for 60 years, less than half the time of FAM-1.

Greater subclonal variation was found in FAM-1 than US-1 genotypes on all continents but Africa, where US-1 was more diverse (Table 1). There were 85 multilocus genotypes (MLGs) of FAM-1 and 36 MLGs of US-1 (Table 1). The greatest diversity of FAM-1 MLGs occurred in specimens from North America, Europe and Asia. FAM-1 displayed higher diversity values across all calculated indices, as well as a higher index of association (Ia) and standardized index of association 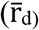 than US-1.

The majority of the herbarium samples (111) were collected from potato, while only 13 were from tomato (Table 1). The remaining specimens were from petunia or wild species of *Solanum species*. There were higher numbers of MLGs from potato than tomato, and greater genetic diversity and higher indices of association/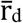 were found in specimens from potato than tomato.

Several SSR loci were useful for distinguishing the FAM-1 genotype from US-1. Diagnostic loci were Pi70 (192/192 in FAM-1 and 189/192 in US-1), PiG11 (160/200 in FAM-1 and 152/156/200 in US-1), PinfSSR2 (173/173 in FAM-1 and 173/177 in US-1), and Pi4B (209/213 in FAM-1 and 213/217 in US-1). The average number of alleles was higher in FAM-1 (5.33) than in US-1(4.5) (Supplementary Table 2).

Structure analysis was done and the optimal K value was two based on results from Structure Harvester. At K=2, the Structure analysis grouped samples into two groups based on genotype (FAM-1 or US-1). FAM-1 was found before US-1 on each continent (Fig. 2). However. At K=3, US-1 genotypes were more homogeneous, while FAM-1 genotypes displayed allelic variation between the two remaining K groups. At K=3, it was noted that FAM-1 genotypes showed more allelic diversity and shifted assignment from one K group to the other over time, beginning around 1911. Both FAM-1 and US-1 occurred in many geographic locations.

**Figure 2:**
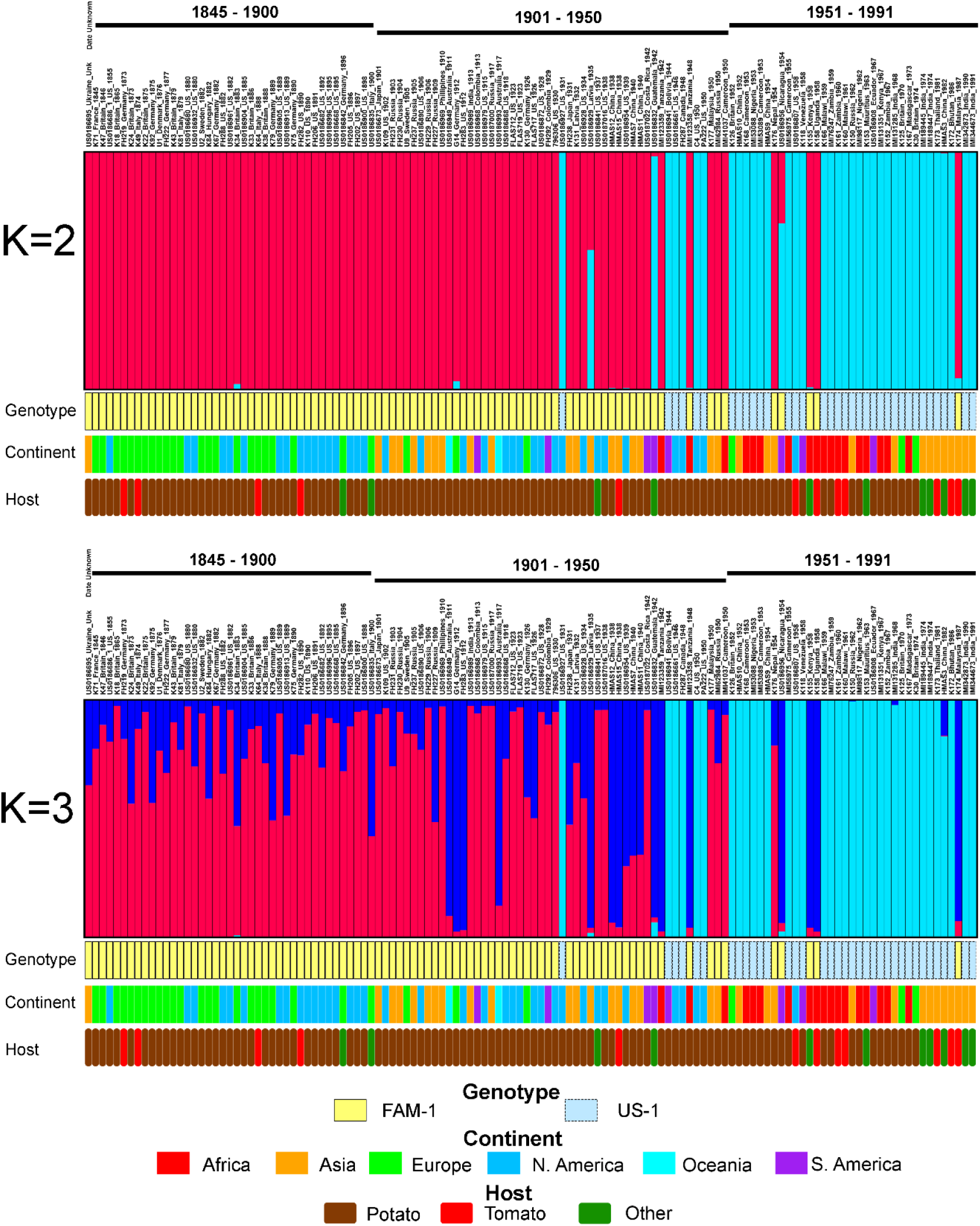
Structure analysis of SSR genotypes of *Phytophthora infestans* from herbarium specimens. Specimens are arranged in ascending chronological order. Samples were clone corrected based on region before analyzing. Results displayed are based under the assumption of two or three groups within the dataset (K=2, K=3). K=2 was calculated to be the most likely based on the second order rate of change.

A minimum spanning network (MSN) was made with haplotypes based on the continent where the samples were collected. Two large groups within the MSN consisting of either FAM-1 or US-1 genotype were observed (Fig. 3). Within the genotype clusters there was no genetic substructuring based on continent. However, the FAM-1 had a larger number of MLGs and more branches in the MSN than the US-1 and greater subclonal variation was observed (Table 1). No exclusive clusters were observed by country, host, or continent, but higher numbers of haplotypes of FAM-1 were found in North America and Asia than elsewhere. There were fewer FAM-1 MLGs from Africa and they occurred on fewer branches of the MSN than MLGs from other continents, most likely due to the more recent introductions there. In contrast, there were more MLGs of US-1 in Africa than North America or Europe, indicating diversification of the US-1 genotype in east Africa.

**Figure 3:**
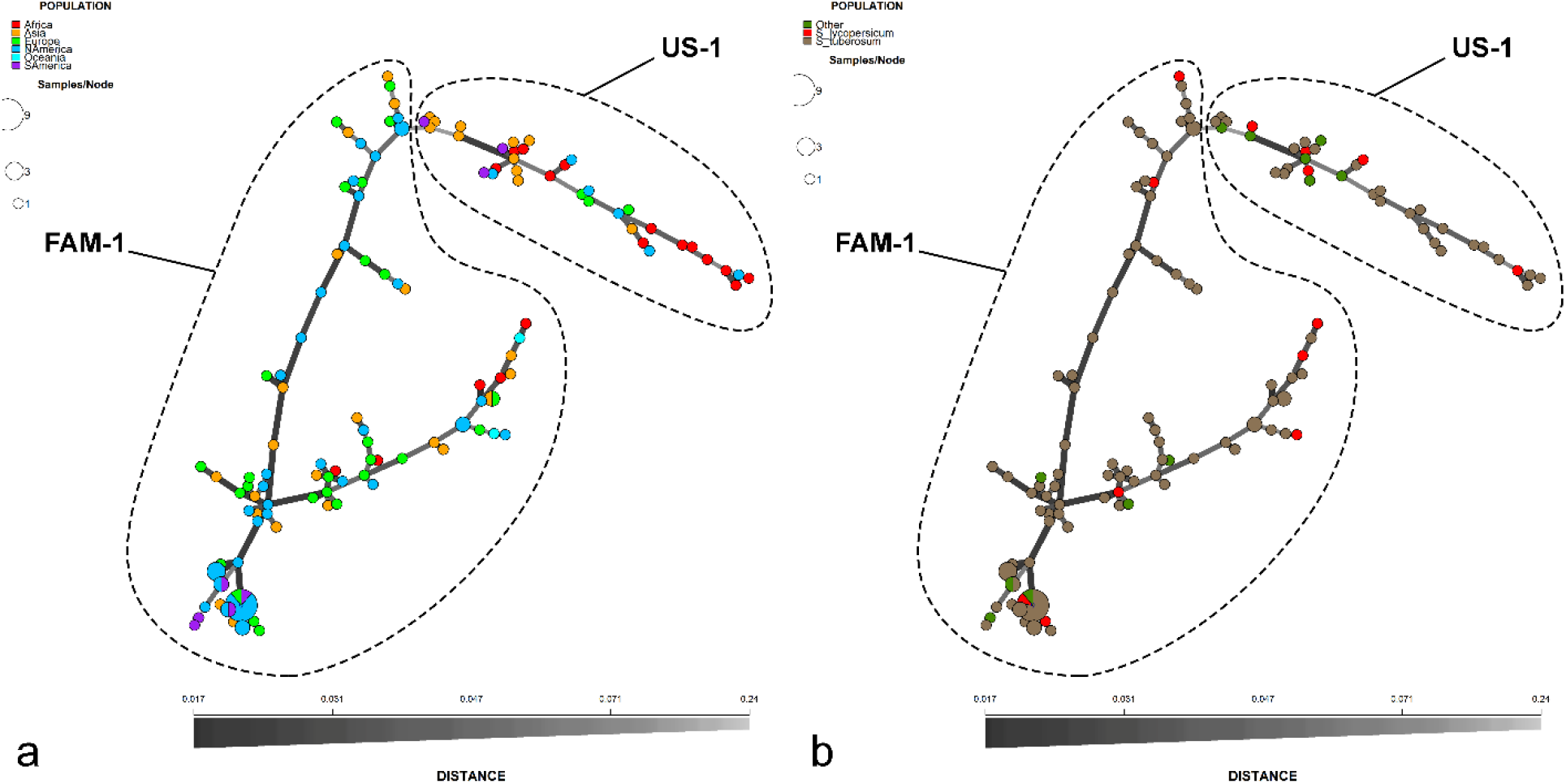
Minimum spanning network of SSR genotypes of *Phytophthora infestans* from herbarium specimens. Data are colored-coded based on (a) the continent where they were collected or (b) host. Genetic distance between haplotypes is indicated by the shade and thickness of the branches.

A similar structure was observed from the neighbor-joining tree, with two large clades that contained either FAM-1 or US-1 genotypes (Fig. 4). When the neighbor-joining tree was expanded to include modern global samples of *P. infestans*, most of the samples identified as FAM-1 or US-1 genotypes also formed homogeneous clusters within the larger neighbor-joining tree using SSRs (Fig. S1).

**Figure 4.**
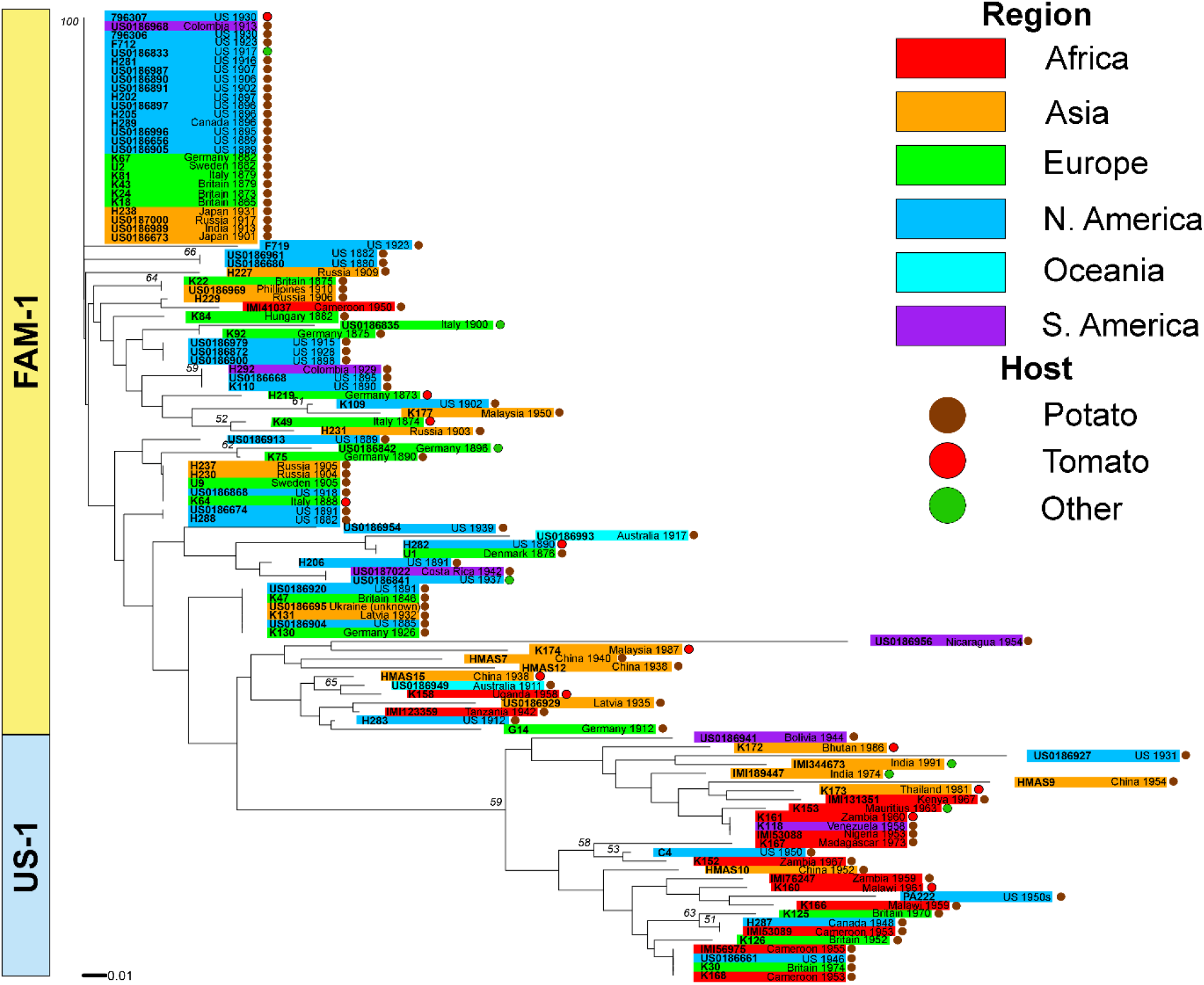
Neighbor joining tree of 7-plex SSR genotypes of *Phytophthora infestan* from herbarium specimens, Specimens are colored-coded based on the continent and host of sample. Bootstrapping was performed with 1000 replicates.

### Migration

Potential migration paths of *P. infestans* into Africa and Asia were tested using specimens identified as FAM-1genotype from Europe and North America. We calculated the probabilities of scenarios that hypothesized movement of *P. infestans* based on a North American source, a European source, and either source with a constant or a varying population size, and a source based on an admixture of European and North American populations (Fig. 5).

**Figure 5.**
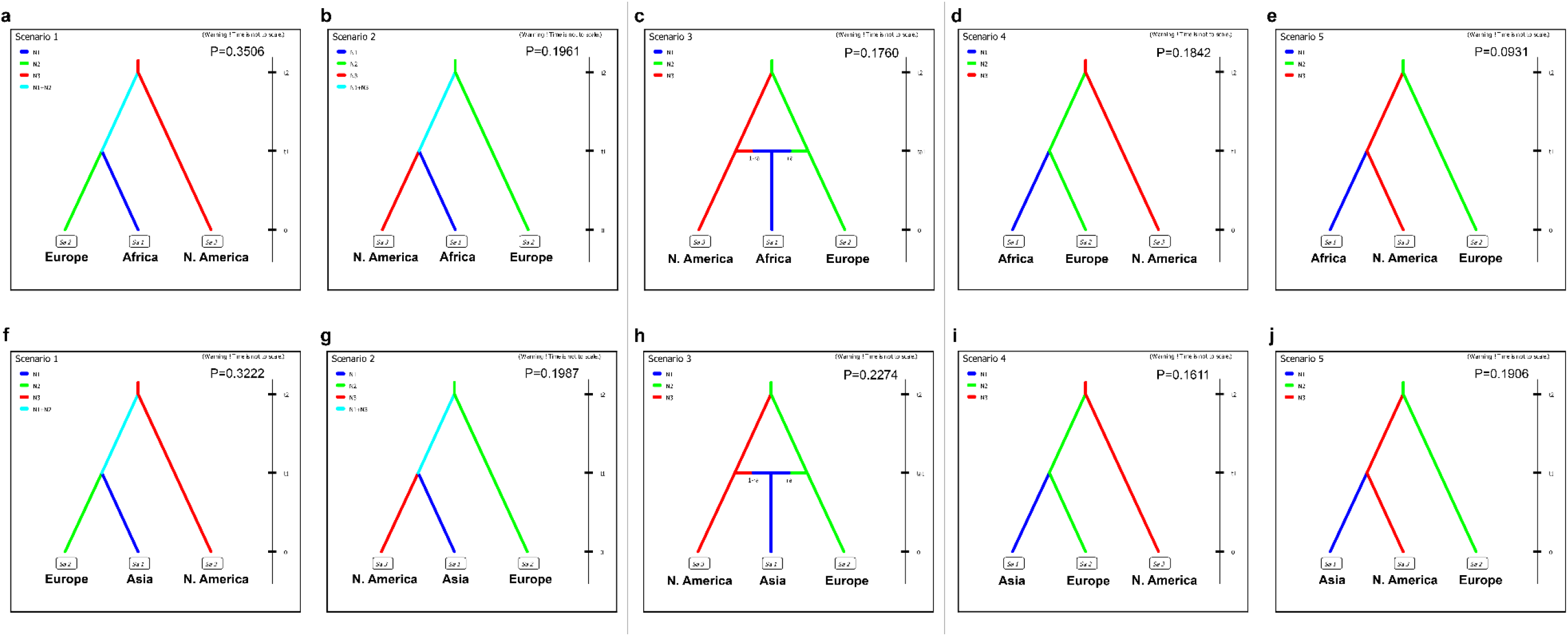
Posterior probabilities of migration scenarios involving populations of the FAM-1 genotype of *Phytophthora infestans* from North America and Europe to Africa(a-e) or North America and Europe to Asia (f-j). Probabilities are based on 1% of the simulated data.

The most likely scenario indicated a divergence first of the European genotypes of FAM-1 from a North American source, followed by a divergence of the African and or Asian FAM-1 genotypes occurred from a European source (Scenario 1, P=0.3506 and 0.3222 for Africa and Asia, respectively) (Supplementary Table 3). Confidence in the scenario choice was evaluated by using simulated datasets to calculate error percentages between the three scenarios with the highest probabilities. Estimation of type I error for scenarios using the Asian data revealed that 53.2% of simulated datasets using this scenario resulted in the highest posterior probability for Scenario 1 when compared to the two scenarios with the next highest probabilities (Scenarios 3, 2) (type I error, 0.468) (Fig. 5 and Supplementary Table 3). For scenarios using the African data, estimation of type I error indicated that 35.6% of simulated datasets resulted in the highest posterior probability for Scenario 1 when compared to the next highest probabilities (Scenarios 2, 4) (type I error, 0.644).

## Discussion

We examined historic outbreaks of *P. infestans* from global historic sources to better understand the history of the spread of the pathogen after the first recorded outbreaks in Europe and the US using herbarium voucher specimens collected worldwide. Our data revealed the widespread presence of the FAM-1 genotype throughout the world and its dominance until the 1930s, when the US-1 lineage began to emerge globally. By the end of the 1950s, FAM-1 had almost completely disappeared from collections and was displaced by US-1 genotype, most likely through movement of potatoes with resistance breeding efforts^2,10^. The only post-1950s FAM-1 genotype observed was collected in a single sample from Malaysia in 1987. This unusual sample suggests that the FAM-1 genotype may have continued to persist in remote areas of the world for a longer period. The Malaysian FAM-1 genotype had variable alleles at several loci when compared to earlier FAM-1 genotypes, suggesting the accumulation of mutations over time or potential hybridization with another lineage such as US-1. It clustered closely with the US-1 genotype in a neighbor- joining tree of a larger set of samples. Further work is underway to sequence the genome of this specimen to understand the variation in more detail.

The earliest known record of the US-1 genotype in potato is from 1931 in the US^10^. Based on specimens analyzed in this study, the earliest known records of US-1 in Africa are from 1953 in Cameroon and Nigeria. For Asia, the oldest US-1 sample observed in this study was from 1952 in China. In South America, the oldest US-1 lineage was from Bolivia in 1944. US-1 was not identified from the samples we examined from Australia in this study, although further work is underway in our lab to genotype more specimens from Australia.

There is scarce information on the history of the early emergence of US-1, but records from the literature suggest an approximate period for multiple parts of the world. In 1947, while documenting the history of late blight in Tasmania, Oldaker commented on an unusual outbreak of *P. infestans* in 1938, in which disease was more sporadic than it had been in previous years, but treatment with copper formulations proved effective in controlling the pathogen^18^. In 1951, Nattrass wrote that potatoes bred for *P. infestans* resistance were failing with the emergence of a new biotype that appeared in Tanzania (then called Tanganyika) in 1946^19^. Our data suggest the new biotype observed was likely the US-1 lineage in Africa. The proximity of these outbreaks suggests that US-1 genotype was spread during potato breeding trials, facilitated by the continued movement of tubers over long distances.

Both the FAM-1 and US-1 genotype predominantly formed two clusters that excluded all modern lineages from Europe, North America, and South America. Our previous studies of North American populations of *P. infestans* suggested a Mexican origin for many of the recent lineages of *P. infestans* circulating in the US^14^. The US-1 genotype has been displaced by other lineages emerging from either Europe or Mexico with metalaxyl resistance or the ability to overcome host resistance genes being bred into potatoes and tomatoes at the time.

Migration analyses of FAM-1 outbreak samples was analyzed using DIYABC analysis and data suggest that both the African and Asian genotypes of FAM-1 most likely emerged from a European source. Outbreaks of the disease caused by FAM-1 genotype first occurred in North America and subsequently spread to Europe^10^. This coincides with historic records and our previous studies that support the migration of *P. infestans* into Europe after outbreaks occurred in North America^2,10,22^. The top migration scenario for emergence into Africa and Asia is from a European source. We do not attempt to identify source countries within Europe but likely sources from historical records include countries such as the UK and or the NL.

Our data and examination of global herbarium sources suggest that the pathogen likely moved first on potatoes and then spread later into tomato^22^. There was greater genetic diversity among potato than tomato genotypes of FAM-1 and more MLGs among the potato genotypes. There were also more infected potato than tomato specimens in a global search of archival collections^22^. The FAM-1 genotype was also more genetically diverse than the US-1 genotype.

The findings of our study are supported by historical reports published by researchers contemporary to the time of the initial outbreaks and provide insight into potential sources^4,17–19,23^. Potatoes were disseminated across the world by European sailors and missionaries, with varying degrees of adoption by local populations^24^. As European colonists moved into new regions, potatoes moved with them. Potatoes were actively encouraged as food for native people during colonization and touted as cheap and nutritious. Potatoes were cited by European scholars to “elevate the happiness and well-being of native peoples”, and subsequently were useful for developing the labor force of the colonizing empire^25^. In India, this mentality resulted in the dissemination of potatoes to local villages by horticultural and agricultural societies, despite the crop having already been adopted as a cash crop to sell to British soldiers^25,26^.

Potatoes continued to move into colonial regions long after their establishment. A colonial handbook for Kenya from 1920 states that potatoes grown from locally produced seed were not as productive, and recommended regularly importing fresh seed potatoes from Europe. This would have provided an obvious avenue for the introduction of *P. infestans*^27^. Regular imports of potatoes were observed in other parts of the world as well. A 1904 agricultural report for the Philippines compared the quality of natively grown potatoes to imported ones found in Manila markets, suggesting a potential introduction route, mostly likely via the US^28^. In India, the pathogen was reported in the area circa 1870-1880 based on reports from local agri-horticultural societies. In letters it was stated that a major late blight outbreak in the Nilgiri region around 1893 was the result of the importation of potatoes from a large nursery in England^17^. In East Africa it was believed that the first outbreak located outside of Nairobi, Kenya, was the result of an importation of Kerr’s Pink potatoes for planting from the United Kingdom, bolstered by a wet and rainy season^4^. In West Africa, however, it was thought that *P. infestans* was introduced as the result of the importation of potatoes from France. While intended for use as food, potatoes were also planted, resulting in the propagation of the pathogen^23^. *Phytophthora infestans* followed the movements by colonists of potato, leading to its introduction from the US and Europe into the African, Asian and Austalian/Oceanic continents.

With the extensive reach of the British Empire (Fig. 1), it is likely that many introductions of the pathogen were the result of movement of potatoes on British ships, with multiple introductions over time as new shipments of tubers were imported into colonies. We are currently doing whole genome analysis of globally sourced specimens to provide more information on the role of host diversity and host jumps in global pathogen spread. The FAM-1 genotype was diploid^11^ and asexual, was able to colonize susceptible potato on six continents and thus caused global pandemics. Our data document that the FAM-1 genotype adapted to many environments, occurred mostly on potato, and remained aggressive for over 140 years.

## Methods

Over 1280 late blight specimens collected in the 19^th^ and 20^th^ century are in herbaria on six continents^22^. We sampled specimens from 37 countries on six continents including North America, South America, Europe, Africa, Asia and Oceania (Fig. 1) (Supplementary Table 1). *Phytophthora infestans* was sampled from herbarium specimens collected in Africa, Asia, Europe, Oceania, North America, and South America. A total of 137 samples were genotyped with microsatellites, consisting of 18 African samples (1942 – 1973), 32 Asian samples (1901 – 1991), 2 Oceania samples (1911 – 1917), 29 European samples (1873 – 1970), 48 North American samples (1855 – 1958), and 8 South American samples (1913 – 1967) (Supplementary Table 1).

We collected SSR data for an additional 194 samples was obtained from databases, published studies, and theses^10,29,30^. These included data from modern populations from Saville et al.^10^ and representatives of current common European lineages from the outbreak tracking system Euroblight.org. In addition we included a subset of data from publications on current Mexican populations^29,30^.

### DNA Extraction, PCR, and Genotyping

DNA was extracted from lesions present on each herbarium voucher using either a Qiagen DNEasy Plant Mini Kit (Qiagen, Valencia, CA) or a modified CTAB method using DNEasy Plant Mini Kit spin columns for cleaning and purifying DNA^12^. The presence of *P. infestans* DNA was checked using species specific primers^31^. All work with herbarium DNA was conducted in a lab in which no modern DNA of *P. infestans* is used, using separate equipment and reagents.

Mitochondrial haplotyping of samples was conducted using primers and PCR cycling conditions developed by Griffith and Shaw to detect the presence of the Ib haplotype^11^,^32^. For detecting the Herb-1 haplotype we utilized primers previously developed that target a single nucleotide polymorphism within the haplotype^10^. Amplicons were sequenced at the Genomic Sciences Laboratory at North Carolina State University.

Samples were genotyped using a 12-plex system of primers for the identification of *P. infestans* lineages using microsatellite loci^33^. To compensate for the low levels of DNA present in extractions, a modification of the PCR protocol was used that increased primer concentration, sample size, and cycling times^15^. The Qiagen Type-It Microsatellite PCR kit (Qiagen Corporation, Valenica CA) was used for PCR reactions, and sample volumes were modified to run a 12.5μL reaction, consisting of 6.25μl of Type-It 2X master mix, 1.3μl of a 10X primer mix (Supplementary Table 4), 1.95μl ddH2O, and 1 – 3μl of DNA extract. Thermal cycling conditions consisted of initial denaturation at 95°C for 5 min, followed by 33 cycles of 95°C for 30 seconds, 58°C for 90 seconds, and 72°C for 30 seconds, and then a final extension period for 30 minutes at 60°C. Fragments were analyzed on an Applied Biosystems 3730xl DNA analyzer at the Genomic Sciences Laboratory at North Carolina State University using 1-3μl of PCR product in a 10.3μL reaction mix consisting of 10μL highly deionized formamide and 0.3μL LIZ500 size standard (Applied Biosystems, Foster City, CA).. Alleles were scored in Geneious 11.1.5 (Biomatters Ltd., Auckland, NZ) using microsatellite plugin 1.4.6. Alleles were named using bin ranges from previously published work^33,34^.

### SSR Data Analysis

Because of the age of herbarium DNA, recovery rates of microsatellite loci from *P. infestans* are lower than they would be for DNA extracted from modern samples, resulting in increased missing data. To reduce the amount of variability due to missing data, only samples with data from at least five SSR loci were used. Data were divided into six categories based on continent: Africa (Afr), Asia (As), Europe (EU), North America (NA), Oceania (Oc), and South America (SA). The broad structure of the populations was evaluated via model-based Bayesian clustering using the program Structure v. 2.3.3 ^35^. Before analysis by Structure, the data were clone corrected (clones were removed such that each population contains only one representative of each haplotype) using the R library poppr v. 2.8.1 ^36^ and R v. 3.5.2 ^37^. Data were clone corrected using their region of collection as a population. The data were run using a 20,000 repeat burn-in and 1,000,000 MCMC repeats under a no admixture model, with each individual sample representing its own population. Independent runs of the model used *K* values from 1 to 10 with 10 replicate runs at each value of *K*. The optimal *K* was estimated using the second order rate of change (the “Evanno method”) in the web tool Structure Harvester ^38,39^. All runs for the optimal *K* value, as well as non-optimal *K* values, were averaged using CLUMPP v. 1.1.2 ^40^ using the Greedy algorithm (M=2) with the pairwise matrix similarity statistic G’ (S=2). The Greedy algorithm was used with 1000 repeats of randomly selected input orders. The resulting output was visualized with the program distruct v. 1.1^41^. Poppr was also used to infer population statistics including: the number of samples (N), the number of multilocus genotypes (MLG), the number of expected MLGs at the smallest sample size of at least 10 (eMLG)^42^, the Shannon-Weiner index of MLG diversity (H)^43^, the Stoddart and Taylor index of MLG diversity (G)^44^, Simpson index corrected for sample size by multiplying the index value by N/(N-1) (λ)^45,46^, evenness^(47–49^, Nei’s unbiased gene diversity (Hexp)^50^, the index of association (Ia)^51,52^, and the standardized index of association 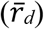^53^.

Relationships between locations and haplotypes of samples were further explored using a minimum spanning network (MSN) based on Bruvo’s distance using the R library adegenet v 2.1.1 ^54^. In addition, a neighbor-joining (NJ) tree based on Bruvo’s distance was constructed using the poppr R library and a combination (genome addition and genome loss) model. In order to utilize a complete dataset in the NJ tree for the purposes of bootstrapping, five loci with low recovery rates were removed for tree construction (PinfSSR8, PinfSSR4, Pi63, PinfSSR11, Pi4B). Any remaining samples still containing missing data were removed. The tree was bootstrapped using 1000 samplings.

An additional 7-plex neighbor joining tree was generated as above using the combined dataset from herbarium and modern samples. Due to the bootstrapping function used (bruvo.boot), no samples with missing data or null alleles could be used. Therefore, samples with putative null alleles were also removed from the dataset for the neighbor-joining tree.

## Migration of FAM-1

Migration routes of the FAM-1 lineage of *P. infestans* into Africa and Asia from Europe and/or North America were examined using Approximate Bayesian Comparison (ABC), as implemented in the program DIYABC v. 2.0.4^55^. Tested migration scenarios for both African and Asian populations included direct divergence from Europe or North America, divergence including a change in population size, or admixture between European and North American populations. Parameter range priors were initialized with values from Saville et al.^10^ and then iteratively modified to better fit our data (Supplementary Table 5). A total of 5 million datasets were simulated. Scenario probabilities were determined through comparison of the observed dataset to simulated datasets generated by DIYABC. A logistic regression of these differences was computed using ten proportions of the simulated dataset as the dependent variable and corresponding differences between the observed and simulated datasets as the independent variable. The value calculated using 50,000 simulated datasets was taken as the scenario’s overall probability. Confidence in the highest scenario was evaluated using a type I error tests, in which the data were compared against 500 simulated data sets assuming the scenario with the highest probability is true and the number of times the scenario in question was correctly or incorrectly applied to the data was determined.

## Acknowledgments

Appreciation is expressed to the following herbaria for providing herbarium material: Royal Botanic Gardens Kew Mycological Herbarium (K), the National Botanic Garden, Glasnevin, Dublin (DBN), the USDA National Fungus Collection, Beltsville, MD (BPI), the CABI Bioscience collection, Egham (IMI), the Farlow Herbarium Harvard University, Cambridge, MA (FH), the Museum of Evolutionary Biology, Uppsala University, Uppsala (UPS), the Cornell Plant Pathology Herbarium, Ithaca, NY (CUP), and the University of Florida Herbarium, Gainesville, FL (FLAS). Thanks to David Cooke, James Hutton Institute, for providing a set of European lineages from EuroBlight database shown in Supplemental Figure S1. Funding was provided by USDA AFRI Grant Number 5197-NCSU-USDA-3179 and USDA AFRI Grant 2011-68004-30154 and the North Carolina Agricultural Research Service.

## Author Contribution

Jbr collected the samples and conceived experiments. AS conducted experiments and analysed data. JBR and AS interpreted data and co-wrote the paper. JBR and AS contributed equally as coauthors of this work

## Data Availability

Raw SSR data and binning rules can be found on GitHub link to be provided once paper is accepted. See the references in the Supplementary Information for data used in the analysis.

## Notes

### Competing Interest Statement

The authors have declared no competing interest.

