## Supplementary Tables and Figures for "Global historic pandemics caused by the FAM-1 genotype of the Irish potato pathogen *Phytophthora infestans*"

**Supplementary Table 1**. Sample number, date of collection, location, genotype or mitochondrial lineage, host and collector of isolates and herbarium specimens of *Phytophthora infestans* used in this study.

| Sample Number | Date | Country | Continent | Genotype/ mtDNA haplotype | Host | Collector/Source |
|---|---|---|---|---|---|---|
| **HERBARIUM** | | | | | | |
| IMI 123359 | 1942 | Tanzania | Africa | FAM-1/Herb-1 | Potato | Unknown |
| IMI 123358 | 1948 | Tanzania | Africa | FAM-1/Not Ib | Potato | Unknown |
| IMI 41037 | 1950 | Cameroon | Africa | FAM-1/Herb-1 | Potato | S. Johnson |
| IMI 43644 | 1950 | Kenya | Africa | ND | *Solanum incanum* | R. Nattrass |
| K 168 | 1953 | Cameroon | Africa | US-1/Ib | Potato | J.A. Russell |
| IMI 53088 | 1953 | Nigeria | Africa | US-1/Ib | Potato | T.A. Russell |
| IMI53089 | 1953 | Cameroon | Africa | US-1/Ib | Potato | T.A. Russell |
| BPI US0186922 | 1955 | Ethiopia | Africa | ND | Potato | R. Stewart |
| IMI 56975 | 1955 | Cameroon | Africa | US-1/Ib | Potato | T.A. Russell |
| K 155 | 1958 | Kenya | Africa | FAM-1/Herb-1 | *S. incanum* | R. Nattrass |
| K 158 | 1958 | Uganda | Africa | FAM-1/Herb-1 | Tomato | G.G. Williams |
| K 166 | 1959 | Malawi | Africa | US-1/Ib | Potato | D.M. Corbett |
| IMI 76247 | 1959 | Zambia | Africa | US-1/Ib | Potato | D.C. Corbett |
| K 161 | 1960 | Zambia | Africa | US-1/Ib | Tomato | A. Angus |
| K 160 | 1961 | Malawi | Africa | US-1/Ib | Tomato | D. Corbett |

| | | | | | | |
|---|---|---|---|---|---|---|
| IMI 98117 | 1962 | Nigeria | Africa | US-1/Ib | Potato | G.G. Williams |
| K 153 | 1963 | Mauritius | Africa | US-1/Ib | Petunia | L. Orieux |
| K 152 | 1967 | Zambia | Africa | US-1/Ib | Potato | A. Rothwell |
| IMI 131351 | 1967 | Kenya | Africa | US-1/Ib | Potato | M. Mogk |
| K 167 | 1973 | Madagascar | Africa | US-1/Ib | Potato | L. Sundheim |
| BPI US0186673 | 1901 | Japan | Asia | FAM-1/Herb-1 | Potato | Fukahashi |
| FH 231 | 1903 | Russia | Asia | FAM-1/Herb-1 | Potato | F. Bucholtz |
| FH 230 | 1904 | Russia | Asia | FAM-1/Herb-1 | Potato | F. Bucholtz |
| FH 237 | 1905 | Russia | Asia | FAM-1/ND | Potato | F. Bucholtz |
| FH 229 | 1906 | Russia | Asia | FAM-1/ND | Potato | F. Bucholtz |
| FH 227 | 1909 | Russia | Asia | FAM-1/ND | Potato | F. Bucholtz |
| BPI US0186969 | 1910 | Philippines | Asia | FAM-1/Herb-1 | Potato | H.S. Yates |
| BPI US0186989 | 1913 | India | Asia | FAM-1/Herb-1 | Potato | J.F. Dastur |
| BPI US0187000 | 1913 | Russia | Asia | FAM-1/ND | Potato | F. Buchoitz, A. Bondarzow |
| FH 238 | 1931 | Japan | Asia | FAM-1/Herb-1 | Potato | K. Togashi |
| K 131 | 1932 | Latvia | Asia | FAM-1/Herb-1 | Potato | K. Starcs |
| BPI US0186929 | 1935 | Latvia | Asia | FAM-1/Herb-1 | Potato | K. Starcs |
| HMAS 12 | 1938 | China | Asia | FAM-1/Herb-1 | Potato | C.C. Cheo |
| HMAS 15 | 1938 | China | Asia | FAM-1/ND | Tomato | F.L. Tai |
| HMAS 7 | 1940 | China | Asia | US-1/ND | Potato | Q. Yuan |

| | | | | | | |
|---|---|---|---|---|---|---|
| HMAS 11 | 1940 | China | Asia | FAM-1/ND | Potato | H. Zhang-xun |
| K 177 | 1950 | Malaysia | Asia | FAM-1/Herb-1 | Potato | A. Johnston |
| IMI 62984 | 1950 | Russia | Asia | FAM-1/ND | Potato | Unknown |
| HMAS 10 | 1952 | China | Asia | US-1/Ib | Potato | Y. Zuo-min |
| HMAS 9 | 1954 | China | Asia | US-1/Ib | Potato | H. He |
| K 176 | 1954 | Nepal | Asia | FAM-1/Herb-1 | Potato | Staunton |
| K 150 | 1962 | Russia | Asia | US-1/Not Herb-1 | Potato | E. Ljegenjakaja |
| IMI 131285 | 1968 | India | Asia | US-1/Ib | Potato | J.A. Russell |
| IMI 189445 | 1974 | India | Asia | US-1/Ib | *S. laciniatum* | D.N. Bordoloi |
| IMI 189447 | 1974 | India | Asia | US-1/Ib | *S. marginatum* | D.N. Bordoloi |
| K 173 | 1981 | Thailand | Asia | US-1/Ib | Tomato | R. Black |
| HMAS 3 | 1982 | China | Asia | US-1/Ib | *S. lyratum* | Q. Yun |
| K 172 | 1986 | Bhutan | Asia | US-1/Ib | Tomato | W.T.D. Peregonne |
| K 174 | 1987 | Malaysia | Asia | FAM-1/Herb-1 | Tomato | B.C. Sutton |
| IMI 342873 | 1990 | India | Asia | US-1/Ib | *S. xanthocarpum* | G.S. Hall |
| IMI 344673 | 1991 | India | Asia | US-1/Ib | *S. meloangena* | G.S. Hall |
| BPI US0186695 | Unk. | Ukraine | Asia | FAM-1/Herb-1 | Potato | R. Tolf |
| BPI US0186949 | 1911 | Australia | Australia/ Oceania | FAM-1/Herb-1 | Potato | B. McAlpine |
| BPI US0186993 | 1917 | Australia | Australia/ Oceania | FAM-1/Herb-1 | Potato | W.A. Birmingham |
| K 71 | 1845 | France | Europe | FAM-1/ND | Potato | J. B. Desmazieres |

| | | | | | | |
|---|---|---|---|---|---|---|
| K 47 | 1846 | Britain | Europe | FAM-1/ND | Potato | M.J. Berkeley |
| K18 | 1865 | Britain | Europe | FAM-1/ND | Potato | M.C. Cooke |
| K 24 | 1873 | Britain | Europe | FAM-1/Herb-1 | Potato | J. E. Vize |
| FH 219 | 1873 | Germany | Europe | FAM-1/Herb-1 | Tomato | F. von Thümen |
| K 49 | 1874 | Italy | Europe | FAM-1/Herb-1 | Tomato | P.A. Saccardo |
| K 22 | 1875 | Britain | Europe | FAM-1/Herb-1 | Potato | J.E. Vize |
| K 92 | 1875 | Germany | Europe | FAM-1/ND | Potato | de Thumen |
| UPS 1 | 1876 | Denmark | Europe | FAM-1/Herb-1 | Potato | E. Rostrup |
| FH 222 | 1877 | Germany | Europe | FAM-1/Herb-1 | Potato | P. Magnus |
| K 43 | 1879 | Britain | Europe | FAM-1/Not Ib | Potato | M.J. Berkeley |
| K 81 | 1879 | Italy | Europe | FAM-1/Herb-1 | Potato | C. Spegazzini |
| K 67 | 1882 | Germany | Europe | FAM-1/ND | Potato | P. Sydow |
| UPS 2 | 1882 | Sweden | Europe | FAM-1/Herb-1 | Potato | J. Eriksson |
| K 84 | 1882 | Hungary | Europe | FAM-1/Not Ib | Potato | G. Linhart |
| K 34 | 1883 | Britain | Europe | FAM-1/Herb-1 | Potato | W.B. Grove |
| K 33 | 1886 | Britain | Europe | FAM-1/ND | Potato | J.E. Vize |
| K 38 | 1888 | Britain | Europe | FAM-1/Herb-1 | Potato | M. Cooke |
| K 64 | 1888 | Italy | Europe | FAM-1/Herb-1 | Tomato | G. Cavara, F. Briosi |
| K 79 | 1889 | Germany | Europe | FAM-1/ND | Potato | P. Hennings |
| K 75 | 1890 | Germany | Europe | FAM-1/ND | Potato | W. Krieger |

| | | | | | | |
|---|---|---|---|---|---|---|
| BPI US0186842 | 1896 | Germany | Europe | FAM-1/ND | *S. nigrum* | P. Sydow |
| BPI US0186835 | 1900 | Italy | Europe | FAM-1/ND | *S. dulcamara* | T. Ferraris |
| UPS 9 | 1905 | Sweden | Europe | FAM-1/Herb-1 | Potato | J. Eriksson |
| G 14 | 1912 | Germany | Europe | FAM-1/ND | Potato | H. Zimmerman |
| K 130 | 1926 | Germany | Europe | FAM-1/ND | Potato | Eliasson |
| K 126 | 1952 | Britain | Europe | US-1/Ib | Potato | J.H.H. |
| K 125 | 1970 | Britain | Europe | US-1/Ib | Potato | R.W. Dennis |
| K 30 | 1974 | Britain | Europe | US-1/Ib | Potato | R.W. Dennis |
| BPI US0186686 | 1855 | US (NY) | N. America | FAM-1/ND | Potato | J.B. Ellis |
| BPI US0186680 | 1880 | US (WI) | N. America | FAM-1/Herb-1 | Potato | W. Trelease |
| BPI US0186932 | 1880 | US (ME) | N. America | FAM-1/Herb-1 | Potato | F.L. Harvey |
| FH 288 | 1882 | US (IL) | N. America | FAM-1/Herb-1 | Potato | A.B. Seymour |
| BPI US0186961 | 1882 | US (WI) | N. America | FAM-1/Herb-1 | Potato | G.A. Pabodie |
| BPI US0186904 | 1885 | US (NY) | N. America | FAM-1/Herb-1 | Potato | W.R. Dudley |
| BPI US0186905 | 1889 | US (NY) | N. America | FAM-1/Herb-1 | Potato | W.R. Dudley |
| BPI US0186656 | 1889 | US (MA) | N. America | FAM-1/Herb-1 | Potato | W.C. Sturgis |
| BPI US0186913 | 1889 | US (MA) | N. America | FAM-1/Herb-1 | Potato | J.E. Humphrey |
| FH 282 | 1890 | US (CT) | N. America | FAM-1/ND | Tomato | R. Thaxter |
| K 110 | 1890 | US (TN) | N. America | FAM-1/ND | Potato | F.L. Scribner |
| FH 206 | 1891 | US (NC) | N. America | FAM-1/Herb-1 | Potato | A.B. Seymour |

| | | | | | | |
|---|---|---|---|---|---|---|
| BPI US0186674 | 1891 | US (MD) | N. America | FAM-1/Herb-1 | Potato | W.T. Swingle |
| BPI US0186920 | 1891 | US (NY) | N. America | FAM-1/Herb-1 | Potato | M.B. Thomas |
| BPI US0186996 | 1891 | US (VT) | N. America | FAM-1/Herb-1 | Potato | L.R. Jones |
| BPI US0186682 | 1892 | US (VT) | N. America | FAM-1/ND | Potato | L.R. Jones |
| BPI US0186668 | 1895 | US (TN) | N. America | FAM-1/Herb-1 | Potato | F.L. Scribner |
| FH 289 | 1896 | Canada | N. America | FAM-1/Herb-1 | Potato | J. Dearness |
| FH 205 | 1896 | US (NY) | N. America | FAM-1/Herb-1 | Potato | B.M. Duggar |
| FH 202 | 1897 | US (VT) | N. America | FAM-1/Herb-1 | Potato | F.L. Sargent |
| BPI US0186897 | 1896 | US (VT) | N. America | FAM-1/Herb-1 | Potato | L.R. Jones |
| BPI US0186900 | 1898 | US (VT) | N. America | FAM-1/Herb-1 | Potato | L.R. Jones, Orton |
| K 109 | 1902 | US (VT) | N. America | FAM-1/ND | Potato | G.P. Clinton |
| BPI US0186891 | 1902 | US (CT) | N. America | FAM-1/Herb-1 | Potato | G.P. Clinton |
| BPI US0186882 | 1902 | US (CT) | N. America | FAM-1/Herb-1 | Potato | G.P. Clinton |
| BPI US0186890 | 1906 | US (CT) | N. America | FAM-1/Herb-1 | Potato | G.P. Clinton |
| BPI US0186987 | 1907 | US (ME) | N. America | FAM-1/Herb-1 | Potato | L.R. Jones |
| FH 283 | 1912 | US (NY) | N. America | FAM-1/Herb-1 | Potato | C. Chupp |
| BPI US0186979 | 1915 | US (PA) | N. America | FAM-1/ND | Potato | G. R. Lyman |
| FH 281 | 1916 | US (ME) | N. America | FAM-1/Herb-1 | Potato | R. Thaxter |
| BPI US0186833 | 1917 | US (DC) | N. America | FAM-1/ND | *Solanum sp.* | W.H. Weston |
| BPI US0186868 | 1918 | US (CT) | N. America | FAM-1/Herb-1 | Potato | G.P. Clinton |

| | | | | | | |
|---|---|---|---|---|---|---|
| FLAS 712 | 1923 | US (FL) | N. America | FAM-1/Herb-1 | Potato | Weber |
| FLAS 719 | 1923 | US (FL) | N. America | FAM-1/ND | Potato | Weber |
| FLAS 717 | 1926 | US (FL) | N. America | FAM-1/ND | Potato | Weber |
| BPI US0186872 | 1928 | US (CT) | N. America | FAM-1/Herb-1 | Potato | G.P. Clinton |
| BPI 796306 | 1930 | US (WV) | N. America | FAM-1/Herb-1 | Potato | W.A. Orton |
| BPI 796307 | 1930 | US (WV) | N. America | FAM-1/Herb-1 | Tomato | W.A. Orton |
| BPI US0186927 | 1931 | US (TX) | N. America | US-1/Ib | Potato | W.J. Bach |
| BPI US0186928 | 1934 | US (AK) | N. America | FAM-1/ND | Potato | G.F. Gravatt |
| BPI US0186841 | 1937 | US (OR) | N. America | FAM-1/Herb-1 | *S. nigrum* | R. Sprague |
| BPI US0187016 | 1938 | US (FL) | N. America | FAM-1/ND | Potato | H.A. Edson |
| BPI US0186954 | 1939 | US (WA) | N. America | FAM-1/ND | Potato | L. Campbell |
| BPI US0186661 | 1946 | US (CT) | N. America | US-1/ND | Tomato | A.D. McDonnell |
| FH 287 | 1948 | Canada | N. America | US-1/Ib | Potato | B.O. Saville |
| CUP 4 | 1950 | US (MN) | N. America | US-1/Ib | Potato | D. Thurston |
| PA222 | 1970s | US (PA) | N. America | US-1/Ib | Potato | M. Gallegly |
| BPI US0186807 | 1958 | US (OH) | N. America | US-1/ND | Tomato | J.L. Cunningham |
| BPI US0186968 | 1913 | Colombia | S. America | FAM-1/Herb-1 | Potato | J. M. Vargas Vergara |
| FH 292 | 1929 | Colombia | S. America | FAM-1/Herb-1 | Potato | C. H. Chandon |
| BPI US0187022 | 1942 | Costa Rica | S. America | FAM-1/Herb-1 | Potato | R. Mendez |
| BPI US0186832 | 1942 | Guatemala | S. America | FAM-1/Herb-1 | Petunia | J. A. Stevenson/ Muller |

| | | | | | | |
|---|---|---|---|---|---|---|
| BPI US0186941 | 1944 | Bolivia | S. America | US-1/Ib | Potato | M. Cardenas |
| BPI US0186956 | 1954 | Nicaragua | S. America | FAM-1/Ia | Potato | S.C. Litzenberger |
| K118 | 1958 | Venezuela | S. America | US-1/ND | Potato | R.W. Dennis |
| BPI US0186908 | 1967 | Ecuador | S. America | US-1/Ib | Potato | V.C. Withee |
| **MODERN** | | | | | | |
| 6/95 | 1995 | Ireland | Europe | 8_A1/IIa | Potato | L. Cooke |
| 31/95 | 1995 | Ireland | Europe | 8_A1/IIa | Potato | L. Cooke |
| 3/99 | 1999 | Ireland | Europe | V-3/IIa | Potato | L. Cooke |
| 21A/93 | 1993 | Ireland | Europe | 8_A1/IIa | Potato | L. Cooke |
| 15/99 | 1999 | Ireland | Europe | 8_A1/IIa | Potato | L. Cooke |
| 12/94 | 1994 | Ireland | Europe | ND/Ia | Potato | L. Cooke |
| 16/99 | 1999 | Ireland | Europe | V-1/Ia | Potato | L. Cooke |
| 15/94 | 1994 | Ireland | Europe | IE-3/Ia | Potato | L. Cooke |
| 18/94 | 1994 | Ireland | Europe | 8_A1/IIa | Potato | L. Cooke |
| 94-1 | 1994 | US (NC) | N. America | US-1/Ib | Potato | J. Ristaino |
| 95-6 | 1995 | US (PA) | N. America | US-1/Ib | Potato | B. Christ |
| US920141 | 1992 | US (ND) | N. America | US-1/Ib | Potato | W. Fry |
| 188.1.1 | 1994 | Canada | N. America | US-1/Ib | Potato | Z. Punja |
| 920159 | 1992 | US (WA/OR) | N. America | US-6/IIb | Potato | W. Fry |
| 94-55 | 1994 | US (NY) | N. America | US-6/IIb | Tomato | W. Fry |

| | | | | | | |
|---|---|---|---|---|---|---|
| 94-52 | 1994 | US (NY) | N. America | US-6/IIb | Potato | W. Fry |
| 94-22 | 1994 | US (NC) | N. America | US-7/Ia | Tomato | J. Ristaino |
| 94-11-2 | 1994 | US (NC) | N. America | US-7/Ia | Tomato | J. Ristaino |
| 93-3 | 1993 | US (NC) | N. America | US-7/Ia | Tomato | J. Ristaino |
| 94-53 | 1994 | US (NC) | N. America | US-7/Ia | Potato | J. Ristaino |
| 2.1.3 | 1993 | Canada | N. America | US-7/Ia | Potato | Z. Punja |
| 98-97 | 1998 | US (NC) | N. America | US-8/Ia | Potato | J. Ristaino |
| RS2009P1 | 2009 | US (PA) | N. America | US-8/Ia | Potato | B. Gugino |
| 98-82 | 1998 | US (NC) | N. America | US-8/Ia | Potato | J. Ristaino |
| 94-8-4 | 1994 | US (NC) | N. America | US-8/Ia | Potato | J. Ristaino |
| VA09-pot | 2009 | US (VA) | N. America | US-8/Ia | Potato | J. Ristaino |
| 97-24 | 1997 | US (NC) | N. America | US-8/Ia | Potato | J. Ristaino |
| 342.1.1 | 1995 | Canada | N. America | US-11/IIb | Potato | Z. Punja |
| 268.1.5 | 1995 | Canada | N. America | US-11/IIb | Potato | Z. Punja |
| US980059 | 1998 | US (AK) | N. America | US-11/IIb | Potato | W. Fry |
| US940478 | 1994 | US (WA) | N. America | US-11/IIb | Potato | W. Fry |
| US980042 | 1998 | US (CA) | N. America | US-11/IIb | Tomato | W. Fry |
| TN-070-A | 2007 | US (TN) | N. America | US-22/Ia | Tomato | K. Deahl |
| TNFL-2 | 2007 | US (TN) | N. America | US-22/Ia | Tomato | K. Deahl |
| NY09- | 2009 | US (NY) | N. America | US-22/Ia | Tomato | M. McGrath |

| | | | | | | |
|---|---|---|---|---|---|---|
| BL2009P4 | 2009 | US (PA) | N. America | US-23/Ia | Potato | B. Gugino |
| PSUTomA | 2009 | US (PA) | N. America | US-23/Ia | Tomato | B. Gugino |
| 63EB | 2014 | US (NC) | N. America | US-23/Ia | Tomato | R. Gardner |
| NC870 | 2014 | US (NC) | N. America | US-23/Ia | Tomato | R. Gardner |
| NC1CELBR | 2014 | US (NC) | N. America | US-23/Ia | Tomato | R. Gardner |
| ND884-1 | 2009 | US (ND) | N. America | US-24/Ia | Potato | G. Secor |
| ND888 | 2009 | US (ND) | N. America | US-24/Ia | Potato | G. Secor |
| ND10-936 | 2010 | US (ND) | N. America | US-24/Ia | Potato | G. Secor |
| BOL3 | — | Bolivia | S. America | BR-1/IIa | Potato | PROINPA |
| BOL9 | — | Bolivia | S. America | BR-1/IIa | Potato | PROINPA |
| B217 | 1998 | Brazil | S. America | BR-1/IIa | Potato | E. Mizubuti |
| B219 | 1998 | Brazil | S. America | BR-1/IIa | Potato | E. Mizubuti |
| B189 | 1998 | Brazil | S. America | BR-1/IIa | Potato | E. Mizubuti |
| B193 | 1998 | Brazil | S. America | BR-1/IIa | Potato | E. Mizubuti |
| DR-4 | 2001 | Costa Rica | S. America | CR-1/Ia | Potato | A. Brenes |
| ZB | — | Costa Rica | S. America | CR-1/Ia | Potato | A. Brenes |
| ZE | 2001 | Costa Rica | S. America | CR-1/Ia | Potato | A. Brenes |
| 52 | 2003 | Costa Rica | S. America | CR-1/Ia | *S. longiconicum* | A. Brenes |
| 61 | 2003 | Costa Rica | S. America | CR-1/Ia | Potato | A. Brenes |
| 141 | 2003 | Costa Rica | S. America | CR-1/Ia | Potato | A. Brenes |

| | | | | | | |
|---|---|---|---|---|---|---|
| CI | 2000 | Costa Rica | S. America | CR-1/Ia | Potato | A. Brenes |
| 152 | 2003 | Costa Rica | S. America | CR-1/Ia | Potato | A. Brenes |
| 42 | — | Costa Rica | S. America | CR-1/Ia | Potato | J. Ristaino |
| 92 | — | Costa Rica | S. America | CR-1/Ia | Potato | J. Ristaino |
| 201 | — | Costa Rica | S. America | CR-1/Ia | Potato | J. Ristaino |
| 211 | — | Costa Rica | S. America | CR-1/Ia | Potato | J. Ristaino |
| 71 | — | Costa Rica | S. America | CR-1/Ia | Potato | J. Ristaino |
| 221 | — | Costa Rica | S. America | CR-1/Ia | Potato | J. Ristaino |
| 81 | — | Costa Rica | S. America | CR-1/Ia | Potato | J. Ristaino |
| 231 | — | Costa Rica | S. America | CR-1/Ia | Potato | J. Ristaino |
| 91 | — | Costa Rica | S. America | CR-1/Ia | Potato | J. Ristaino |
| 151 | 2003 | Costa Rica | S. America | CR-1/Ia | Potato | A. Brenes |
| EC3092 | 1997 | Ecuador | S. America | EC-1/IIa | *S. phureja* | G. Forbes |
| EC3094 | 1997 | Ecuador | S. America | EC-1/IIa | *S. phureja* | G. Forbes |
| EC3199 | 1998 | Ecuador | S. America | EC-1/IIa | *S. tuquerense* | G. Forbes |
| EC3253 | 1999 | Ecuador | S. America | EC-1/IIa | *S. columbianum* | G. Forbes |
| EC3154 | 1998 | Ecuador | S. America | EC-1/IIa | *S. andreanum* | G. Forbes |
| EC3298 | 2000 | Ecuador | S. America | EC-1/IIa | *S. tetrapetalum* | G. Forbes |
| EC3300 | 2000 | Ecuador | S. America | EC-1/IIa | *S. paucijugum* | G. Forbes |
| PIC97180 | 1997 | Mexico | S. America | ND/Ia | Potato | N. Grünwald |

| | | | | | | |
|---|---|---|---|---|---|---|
| PIC97630 | 1997 | Mexico | S. America | ND/Ia | Potato | N. Grünwald |
| PIC98392 | 1998 | Mexico | S. America | ND/Ia | *S. demissum* | N. Grünwald |
| PIC97207 | 1997 | Mexico | S. America | ND/Ia | Potato | N. Grünwald |
| PIC97652 | 1997 | Mexico | S. America | ND/Ia | Potato | N. Grünwald |
| PIC97224 | 1997 | Mexico | S. America | ND/Ia | Potato | N. Grünwald |
| PIC98301 | 1998 | Mexico | S. America | ND/Ia | Potato | N. Grünwald |
| PIC98305 | 1998 | Mexico | S. America | ND/Ia | Potato | N. Grünwald |
| PIC98366 | 1998 | Mexico | S. America | ND/Ia | *S. demissum* | N. Grünwald |
| PIC97388 | 1997 | Mexico | S. America | ND/Ia | Potato | N. Grünwald |
| PIC98369 | 1998 | Mexico | S. America | ND/Ia | *S. demissum* | N. Grünwald |
| PIC97605 | 1997 | Mexico | S. America | ND/Ia | Potato | N. Grünwald |
| PIC98372 | 1998 | Mexico | S. America | ND/Ia | *S. demissum* | N. Grünwald |
| PIC97620 | 1997 | Mexico | S. America | ND/Ia | Potato | N. Grünwald |
| PIC98388 | 1998 | Mexico | S. America | ND/Ia | *S. demissum* | N. Grünwald |
| PIC97022 | 1997 | Mexico | S. America | ND | Potato | N. Grünwald |
| PIC97349 | 1997 | Mexico | S. America | ND/Ia | Potato | N. Grünwald |
| PIC98382 | 1998 | Mexico | S. America | ND/Ia | Potato | N. Grünwald |
| PIC97370 | 1997 | Mexico | S. America | ND/Ia | Potato | N. Grünwald |
| PIC97323 | 1997 | Mexico | S. America | ND/Ia | Potato | N. Grünwald |
| PIC97316 | 1997 | Mexico | S. America | ND/Ia | Potato | N. Grünwald |

| | | | | | | |
|---|---|---|---|---|---|---|
| PIC97229 | 1997 | Mexico | S. America | ND/Ia | Potato | N. Grünwald |
| PCZ026 | 1997 | Peru | S. America | PE-6/IIa | Potato | W. Pérez |
| PPU003 | 1997 | Peru | S. America | EC-1/IIa | Potato | W. Pérez |
| PCZ033 | 1997 | Peru | S. America | EC1.1/IIa | Potato | W. Pérez |
| PER802 | 1985 | Peru | S. America | US-1/Ib | Potato | P. Tooley |
| PCZ098 | 1997 | Peru | S. America | EC1.2/IIa | Potato | R. Nelson |
| PCZ118 | 1997 | Peru | S. America | EC1.2/IIa | Potato | W. Pérez |
| PHU006 | 1996 | Peru | S. America | EC-1/IIa | Potato | R. Morales |
| PER832 | 1986 | Peru | S. America | US-1/Ib | Potato | P. Tooley |
| POX004 | 1997 | Peru | S. America | EC-1/IIa | Potato | R. Nelson |
| PCZ007 | 1997 | Peru | S. America | PE-3/Ia | Potato | W. Pérez |
| PCO038 | 1997 | Peru | S. America | EC-1/IIa | Potato | W. Pérez |
| PCZ050 | 1997 | Peru | S. America | PE-3/Ia | Potato | M. Coca |
| PPA008 | 1998 | Peru | S. America | EC-1/IIa | Potato | E. de la Torre |
| PVM004 | 1998 | Peru | S. America | EC-1/IIa | Potato | W. Pérez |
| PER810 | 2008 | Peru | S. America | US-1/Ib | *S. chiquidenum* | P. Tooley |
| PCA014 | 1999 | Peru | S. America | US-1/Ib | Potato | G. Garry |
| PLL018 | 2000 | Peru | S. America | EC-1/IIa | Potato | A. Salas |
| PER803 | 2008 | Peru | S. America | US-1/Ib | Potato | P. Tooley |
| PCA001 | 1999 | Peru | S. America | PE-3/Ia | Potato | G. Garry |

| | | | | | | |
|---|---|---|---|---|---|---|
| PER809 | 2009 | Peru | S. America | US-1/Ib | *S. plurae* | P. Tooley |
| PAN002 | 1999 | Peru | S. America | EC-1/IIa | *S. urophylum* | A. Salas |
| **Modern Mexico**[a] | | | | | | |
| CHG16 | 2015 | Mexico | S. America | ND | Potato | N. Grünwald |
| CHG10 | 2015 | Mexico | S. America | NDsoybean seed dna extraction | Potato | N. Grünwald |
| CHG12 | 2015 | Mexico | S. America | ND | Potato | N. Grünwald |
| CHG32 | 2015 | Mexico | S. America | ND | Potato | N. Grünwald |
| JFH121 | 2015 | Mexico | S. America | ND | Potato | N. Grünwald |
| JFH135 | 2015 | Mexico | S. America | ND | Potato | N. Grünwald |
| JFH163 | 2015 | Mexico | S. America | ND | Potato | N. Grünwald |
| JFH168 | 2015 | Mexico | S. America | ND | Potato | N. Grünwald |
| JFH3 | 2015 | Mexico | S. America | ND | Potato | N. Grünwald |
| JFH41 | 2015 | Mexico | S. America | ND | Potato | N. Grünwald |
| JFH54 | 2015 | Mexico | S. America | ND | Potato | N. Grünwald |
| SGF37 | 2015 | Mexico | S. America | ND | Potato | N. Grünwald |
| SGF1 | 2015 | Mexico | S. America | ND | Potato | N. Grünwald |
| SGF18 | 2015 | Mexico | S. America | ND | Potato | N. Grünwald |
| SGF21 | 2015 | Mexico | S. America | ND | Potato | N. Grünwald |
| SGF66 | 2015 | Mexico | S. America | ND | Potato | N. Grünwald |

| | | | | | | |
|---|---|---|---|---|---|---|
| SGF78 | 2015 | Mexico | S. America | ND | Potato | N. Grünwald |
| SGF79 | 2015 | Mexico | S. America | ND | Potato | N. Grünwald |
| SGF84 | 2015 | Mexico | S. America | ND | Potato | N. Grünwald |
| T18 | 2015 | Mexico | S. America | ND | Potato | N. Grünwald |
| T24 | 2015 | Mexico | S. America | ND | Potato | N. Grünwald |
| TL10 | 2015 | Mexico | S. America | ND | Potato | N. Grünwald |
| TL13 | 2015 | Mexico | S. America | ND | Potato | N. Grünwald |
| TL3 | 2015 | Mexico | S. America | ND | Potato | N. Grünwald |
| TF4 | 2015 | Mexico | S. America | ND | Potato | N. Grünwald |
| TG10 | 2015 | Mexico | S. America | ND | Potato | N. Grünwald |
| CHC78_16 | 2016 | Mexico | S. America | ND | Potato | N. Grünwald |
| JFH104_16 | 2016 | Mexico | S. America | ND | Potato | N. Grünwald |
| JFH108_16 | 2016 | Mexico | S. America | ND | Potato | N. Grünwald |
| JFH109_16 | 2016 | Mexico | S. America | ND | Potato | N. Grünwald |
| JFH122_16 | 2016 | Mexico | S. America | ND | Potato | N. Grünwald |
| JFH49_16 | 2016 | Mexico | S. America | ND | Potato | N. Grünwald |
| JFH59_16 | 2016 | Mexico | S. America | ND | Potato | N. Grünwald |
| JFH8_16 | 2016 | Mexico | S. America | ND | Potato | N. Grünwald |
| JFH93_16 | 2016 | Mexico | S. America | ND | Potato | N. Grünwald |
| SGF39_16 | 2016 | Mexico | S. America | ND | Potato | N. Grünwald |

| | | | | | | |
|---|---|---|---|---|---|---|
| SGF57 | 2016 | Mexico | S. America | ND | Potato | N. Grünwald |
| **WANG ET AL. 2017[b]** | | | | | | |
| Mich7012 | 2007 | Mexico | S. America | ND | Potato | E. Goss |
| Mich7030 | 2007 | Mexico | S. America | ND | Potato | E. Goss |
| Mich7054 | 2007 | Mexico | S. America | ND | Potato | E. Goss |
| **Modern Europe** | | | | | | |
| NL05015 | Unknown | Netherlands | Europe | 13_A2 | Unknown | D. Cooke |
| NL05246 | Unknown | Netherlands | Europe | 13_A2 | Unknown | D. Cooke |
| NL05353 | Unknown | Netherlands | Europe | 13_A2 | Unknown | D. Cooke |
| NL05557 | Unknown | Netherlands | Europe | 13_A2 | Unknown | D. Cooke |
| NL05592 | Unknown | Netherlands | Europe | 13_A2 | Unknown | D. Cooke |
| NL05869 | Unknown | Netherlands | Europe | 13_A2 | Unknown | D. Cooke |
| NL07063 | Unknown | Netherlands | Europe | 13_A2 | Unknown | D. Cooke |
| NL07548 | Unknown | Netherlands | Europe | 13_A2 | Unknown | D. Cooke |
| NL08419 | Unknown | Netherlands | Europe | 13_A2 | Unknown | D. Cooke |
| 07_5202A | 2007 | Unknown | Unknown | 13_A2 | Unknown | D. Cooke |
| 07_4707A | 2007 | Unknown | Unknown | 13_A2 | Unknown | D. Cooke |
| 07_4734B | 2007 | Unknown | Unknown | 13_A2 | Unknown | D. Cooke |
| 07_4774A | 2007 | Unknown | Unknown | 13_A2 | Unknown | D. Cooke |
| 07_5762B | 2007 | Unknown | Unknown | 13_A2 | Unknown | D. Cooke |

| | | | | | | |
|---|---|---|---|---|---|---|
| 08_6650D | 2008 | Unknown | Unknown | 13_A2 | Unknown | D. Cooke |
| 09PL093 | Unknown | Unknown | Unknown | 13_A2 | Unknown | D. Cooke |
| 25_14AN3_5 | Unknown | Unknown | Unknown | 13_A2 | Unknown | D. Cooke |
| NL98014R | Unknown | Netherlands | Europe | 2_A1 | Unknown | D. Cooke |
| 10_8058A | 2010 | Britain | Europe | 2_A1 | Unknown | D. Cooke |
| 07_5622A | 2007 | Britain | Europe | 2_A1 | Unknown | D. Cooke |
| 06_SS7_05 | 2006 | Unknown | Unknown | 2_A1 | Unknown | D. Cooke |
| AU_059 | Unknown | Australia | Oceania | 2_A1 | Unknown | D. Cooke |
| 10_7814A | 2010 | Britain | Europe | 23_A1 | Unknown | D. Cooke |
| 09_7722B | 2009 | Britain | Europe | 23_A1 | Unknown | D. Cooke |
| 07_5974E | 2007 | Britain | Europe | 23_A1 | Unknown | D. Cooke |
| US110059_US23 | 2011 | US(NY) | N. America | 23_A1 | Tomato | J. Wallace |
| BX10-36_TomA_lesion | Unknown | Unknown | Unknown | 23_A1 | Tomato | D. Cooke |
| petunia_pot1_leafC | Unknown | Unknown | Unknown | 23_A1 | Petunia | D. Cooke |
| MSC_11_0014B | 2011 | Unknown | Unknown | 23_A1 | Unknown | D. Cooke |
| MSC_11_0014D | 2011 | Unknown | Unknown | 23_A1 | Unknown | D. Cooke |
| FR_2012_012_002 | 2012 | France | Europe | 23_A1 | Unknown | D. Cooke |
| FR_2012_012_001 | 2012 | France | Europe | 23_A1 | Unknown | D. Cooke |
| EGY_11_25_Men8_1 | 2011 | Egypt | Africa | 23_A1 | Unknown | D. Cooke |

| | | | | | | |
|---|---|---|---|---|---|---|
| C5 | Unknown | Britain | Europe | 5_A1 | Unknown | D. Cooke |
| C4 | Unknown | Britain | Europe | 5_A1 | Unknown | D. Cooke |
| 2006_3872G | 2006 | Unknown | Unknown | 6_A1 | Unknown | D. Cooke |
| 09_7734D | 2009 | Unknown | Unknown | 6_A1 | Unknown | D. Cooke |
| 2005_12637 | 2005 | Unknown | Unknown | 3_A2 | Unknown | D. Cooke |
| 2006_4180_10 | 2006 | Unknown | Unknown | 1_A1 | Unknown | D. Cooke |
| 2006_4388E | 2006 | Unknown | Unknown | 17_A2 | Unknown | D. Cooke |
| Amiga_Desiree_34_2 | Unknown | Unknown | Unknown | 12_A1 | Potato | D. Cooke |
| 06_SS2_09 | 2006 | Unknown | Unknown | 8_A1 | Unknown | D. Cooke |
| PCL_09_7638B | 2009 | Unknown | Unknown | 8_A1 | Unknown | D. Cooke |
| 2015_12346B | 2015 | Unknown | Unknown | 8_A1 | Unknown | D. Cooke |
| 30_14T5_1 | Unknown | Unknown | Unknown | 8_A1 | Unknown | D. Cooke |
| 2005_14635 | 2005 | Unknown | Unknown | 10_A2 | Unknown | D. Cooke |
| Coff_P10105 | 2002 | US(NE) | N. America | US-1 | Potato | C. Smart |
| 10PL074 | Unknown | Unknown | Unknown | 34_A1 | Unknown | D. Cooke |
| PCL_12_9186A | 2010 | Unknown | Unknown | 35_A2 | Unknown | D. Cooke |
| 2014_Amiga_14003A | 2014 | Unknown | Unknown | 36_A2 | Unknown | D. Cooke |
| Eucablight_2015_B_15012 | 2015 | Unknown | Unknown | 36_A2 | Unknown | D. Cooke |
| 2016_12738A | 2016 | Unknown | Unknown | 37_A2 | Unknown | D. Cooke |
| FRANCE_052 | Unknown | France | Europe | 38_A2 | Unknown | D. Cooke |

| | | | | | | |
|---|---|---|---|---|---|---|
| SI1530 | Unkno wn | Unknown | Unknown | 39_A1 | Unknown | D. Cooke |
| 2017_DK_22 | 2017 | Denmark | Europe | 41_A2 | Unknown | D. Cooke |

[a] Shakya, S. K., Larsen, M. M., Cuenca-Condoy, M. M., Lozoya-Saldaña, H. & Grünwald, N. J. Variation in genetic diversity of *Phytophthora infestans* populations in Mexico from the center of origin outwards. *Plant Disease* **102**, 1534-1540, doi:10.1094/PDIS-11-17-1801-RE (2018).

[b] Wang, J. *et al.* High levels of diversity and population structure in the potato late blight pathogen at the Mexico centre of origin. *Molecular Ecology* **26**, 1091-1107, doi:10.1111/mec.14000 (2017).

**Supplementary Table 2.** Diversity indices for 12 microsatellite loci of *Phytophthora infestans*, divided by genotype and clone corrected.

| Genotype[a] | *n*[b] | 1-D | Hexp | Evenness |
|---|---|---|---|---|
| **FAM-1** | | | | |
| D13 | 14 | 0.702 | 0.707 | 0.586 |
| PinfSSR8 | 4 | 0.482 | 0.489 | 0.644 |
| PinfSSR4 | 10 | 0.655 | 0.660 | 0.621 |
| Pi04 | 2 | 0.499 | 0.502 | 0.998 |
| Pi70 | 3 | 0.050 | 0.050 | 0.360 |
| PinfSSR6 | 6 | 0.513 | 0.516 | 0.653 |
| Pi63 | 5 | 0.202 | 0.204 | 0.437 |
| PiG11 | 10 | 0.569 | 0.573 | 0.696 |
| Pi02 | 4 | 0.065 | 0.066 | 0.351 |
| PinfSSR11 | 2 | 0.500 | 0.505 | 1.000 |
| PinfSSR2 | 1 | NA | NA | NA |
| Pi4B | 3 | 0.509 | 0.513 | 0.926 |
| Mean | 5.33 | 0.395 | 0.399 | 0.661 |
| **US-1** | | | | |
| D13 | 9 | 0.730 | 0.740 | 0.710 |
| PinfSSR8 | 3 | 0.160 | 0.160 | 0.460 |
| PinfSSR4 | 9 | 0.770 | 0.780 | 0.740 |
| Pi04 | 2 | 0.500 | 0.500 | 0.990 |
| Pi70 | 2 | 0.500 | 0.510 | 1.000 |
| PinfSSR6 | 5 | 0.420 | 0.430 | 0.530 |
| Pi63 | 4 | 0.670 | 0.690 | 0.940 |
| PiG11 | 9 | 0.740 | 0.750 | 0.760 |
| Pi02 | 2 | 0.030 | 0.030 | 0.380 |
| PinfSSR11 | 2 | 0.460 | 0.470 | 0.930 |
| PinfSSR2 | 2 | 0.500 | 0.510 | 1.000 |
| Pi4B | 4 | 0.530 | 0.540 | 0.860 |
| Mean | 4.42 | 0.500 | 0.510 | 0.780 |
| **ALL** | | | | |
| D13 | 17 | 0.800 | 0.803 | 0.685 |
| PinfSSR8 | 5 | 0.392 | 0.396 | 0.540 |
| PinfSSR4 | 11 | 0.734 | 0.738 | 0.639 |
| Pi04 | 2 | 0.498 | 0.500 | 0.997 |
| Pi70 | 3 | 0.281 | 0.282 | 0.624 |
| PinfSSR6 | 6 | 0.491 | 0.494 | 0.601 |
| Pi63 | 5 | 0.491 | 0.495 | 0.634 |
| PiG11 | 14 | 0.745 | 0.747 | 0.691 |

| | | | | |
|---|---|---|---|---|
| Pi02 | 5 | 0.055 | 0.055 | 0.324 |
| PinfSSR11 | 2 | 0.495 | 0.498 | 0.990 |
| PinfSSR2 | 2 | 0.259 | 0.260 | 0.655 |
| Pi4B | 5 | 0.618 | 0.621 | 0.864 |
| Mean | 6.417 | 0.488 | 0.491 | 0.687 |

[a] All isolates, including those with missing data (minimum=5) are included.

[b] *n*: number of alleles; 1-D: Simpson index; Hexp: Nei's 1978 expected heterozygosity

**Supplementary Table 3.** Migration scenarios and posterior probabilities and confidence intervals for three groups of FAM-1 populations of historic *Phytophthora infestans*.

| Populations[a] | Scenario | Probability [b] | 95% CI |
|---|---|---|---|
| Asia-EU-NA | **1** | **0.3222** | **[0.3043, 0.3401]** |
| | 2 | 0.1987 | [0.1821, 0.2153] |
| | 3 | 0.2274 | [0.2117, 0.2432] |
| | 4 | 0.1611 | [0.1486, 0.1736] |
| | 5 | 0.0906 | [0.0795, 0.1017] |
| Africa-EU-NA | **1** | **0.3506** | **[0.3303, 0.3710]** |
| | 2 | 0.1961 | [0.1776, 0.2145] |
| | 3 | 0.1760 | [0.1599, 0.1922] |
| | 4 | 0.1842 | [0.1589, 0.2094] |
| | 5 | 0.0931 | [0.0771, 0.1090] |

[a] Asia: Asian historic herbarium samples, FAM-1 genotype (1901-1987); Africa: African historic herbarium samples, FAM-1 genotype (1942-1958); EU: European historic herbarium samples, FAM-1 genotype (1845-1926), NA: North American historic herbarium samples, FAM-1genotype (1855-1939).

[b] Probabilities listed are based on 1% of the simulated data.

**Supplementary Table 4.** Primer sequences for 12-plex microsatellites of *P. infestans*.

| Primer | Dye and sequence (5' – 3') | [a]Volume of 100µM stock |
|---|---|---|
| PiG11F | **NED**-TGCTATTTATCAAGCGTGGG | 6 |
| PiG11R | GTTTCAATCTGCAGCCGTAAGA | 6 |
| Pi02F | **NED**-ACTTGCAGAACTACCGCCC | 6 |
| Pi02R | GTTTGACCACTTTCCTCGGTTC | 6 |
| PinfSSR11F | **NED**-TTAAGCCACGACATGAGCTG | 6 |
| PinfSSR11R | GTTTAGACAATTGTTTTGTGGTCGC | 6 |
| D13F | **FAM**-TGCCCCCTGCTCACTC | 6.4 |
| D13R | GCTCGAATTCATTTTACAGACTTG | 6.4 |
| PinfSSR8F | **FAM**-AATCTGATCGCAACTGAGGG | 12 |
| PinfSSR8R | GTTTACAAGATACACACGTCGCTCC | 12 |
| PinfSSR4F | **FAM**-TCTTGTTCGAGTATGCGACG | 6 |
| PinfSSR4R | GTTTCACTTCGGGAGAAAGGCTTC | 6 |
| Pi04F | **VIC**-AGCGGCTTACCGATGG | 6 |
| Pi04R | GTTTCAGCGGCTGTTTCGAC | 6 |
| Pi70F | **VIC**-ATGAAAATACGTCAATGCTCG | 6 |
| Pi70R | CGTTGGATATTTCTATTTCTTCG | 6 |
| PinfSSR6F | **VIC**-GTTTTGGTGGGGCTGAAGTTTT | 6 |
| PinfSSR6R | TCGCCACAAGATTTATTCCG | 6 |
| Pi63F | **VIC**-ATGACGAAGATGAAAGTGAGG | 6 |
| Pi63R | CGTATTTTCCTGTTTATCTAACACC | 6 |
| PinfSSR2F | **PET**-CGACTTCTACATCAACCGGC | 6 |
| PinfSSR2R | GTTTGCTTGGACTGCGTCTTTAGC | 6 |
| Pi4BF | **PET**-AAAATAAAGCCTTTGGTTCA | 12 |
| Pi4BR | GCAAGCGAGGTTTGTAGATT | 12 |
| 10 mM Tris pH 8 | | 168.8 |

[a]Measurements for multiplexing the primers into a 10X master mix are included.

**Supplementary Table 5.** Prior distributions for Do It Yourself Approximate Bayesian Computation (DIYABC) scenarios for FAM-1 historic populations of *Phytophthora infestans*. Summary statistics for all runs: within population statistics: mean number of alleles and mean genetic diversity; between sample statistics: mean number of alleles, mean genetic diversity, Fst, shared allele distance, $(\delta\mu)^2$, and maximum likelihood coefficient of admixture.

| Parameter | Shape | Min — Max |
|---|---|---|
| **Population size** | | |
| Africa FAM-1 | Uniform | 10000 — 400000 |
| Asia FAM-1 | Uniform | 10000 — 400000 |
| Europe FAM-1 | Uniform | 10000 — 500000 |
| North America FAM-1 | Uniform | 10000 — 700000 |
| **Time since divergence**[a] | | |
| t1: Two populations | Uniform | 100 — 20000 |
| t2: Two or three populations | Uniform | 100 — 30000 |
| **Admixture events** | | |
| Admixture rate | Uniform | 0.001 — 0.999 |
| ta1: Timing since admixture event | Uniform | 100 — 20000 |
| **Nucleotide sequence evolution**[b] | | |
| Mean/individual mutation rate | Uniform | $1.00x10^{-8}/1.00\ x10^{-12}$ — $1.00x10^{-7}/1.00x10^{-12}$ |

[a] In order to guide the construction of trees, priors related to the time since divergence and admixture events were defined, such that t2>ta1, t2>t1, and ta1>=t1
[b] The stepwise mutation model was used as the basis of the substitution model

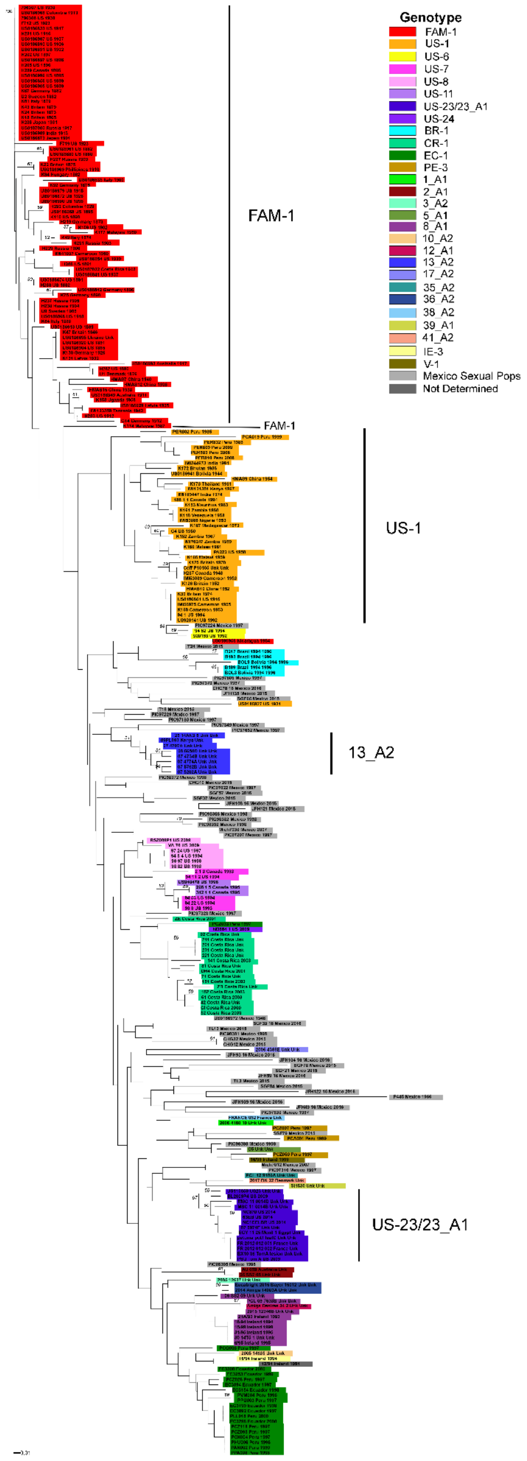

**Supplementary Figure 1**: Neighbor-joining tree 7-plex SSR genotypes of modern and historic populations of *Phytophthora infestans*. Specimens are color-coded based on genotype, if known. Bootstrapping was based on 1000 replicates.
